# Cdc48 targets INQ-localized Mrc1 to facilitate recovery from replication stress

**DOI:** 10.1101/2021.09.16.460581

**Authors:** Camilla Colding, Jacob Autzen, Boris Pfander, Michael Lisby

## Abstract

DNA replication stress is a source of genome instability and a replication checkpoint has evolved to enable fork stabilisation and completion of replication during stress. Mediator of the replication checkpoint 1 (Mrc1) is the primary mediator of this response in *Saccharomyces cerevisiae*. Mrc1 is partially sequestered in the intranuclear quality control compartment (INQ) upon methyl methanesulfonate (MMS)-induced replication stress. Here we show that Mrc1 re-localizes from the replication fork to INQ during replication stress. Sequestration of Mrc1 in INQ is facilitated by the Btn2 chaperone and the Cdc48 segregase is required to release Mrc1 from INQ during recovery from replication stress. Consistently, we show that Cdc48 colocalizes with Mrc1 in INQ and we find that Mrc1 is recognized by the Cdc48 cofactors Ufd1 and Otu1, which contribute to clearance of Mrc1 from INQ. Our findings suggest that INQ localization of Mrc1 and Cdc48 function to facilitate replication stress recovery by transiently sequestering the replication checkpoint mediator Mrc1 and explains our observation that Btn2 and Cdc48 are required for efficient replication restart following MMS-induced replication stress.

## Introduction

Faithful replication of the DNA is essential for cell survival and the transmission of genetic information to succeeding generations. Impediments to DNA replication such as DNA lesions or depletion of deoxynucleotides result in replication fork stalling and checkpoint activation. Failure to activate the checkpoint upon replication stress causes genome instability and is a driver of cancer in humans (Abbas *et al*, 2013). Activation of the replication checkpoint response is conserved from yeast to humans and is initiated by recognition of replication protein A (RPA)-coated single-stranded DNA (ssDNA) resulting from uncoupling of the mini chromosome maintenance (MCM) helicase from the replicative polymerase (Labib & De Piccoli, 2011). In *Saccharomyces cerevisiae*, this recognition is accomplished by Ddc2, which interacts with and recruits the replication checkpoint sensor kinase Mec1 to RPA-coated ssDNA. Replication checkpoint activation is achieved through Mec1-mediated phosphorylation of the checkpoint effector kinase Rad53 at (S/T)Q residues. This is facilitated by mediator of the replication checkpoint 1 (Mrc1), which is also phosphorylated by Mec1 and Rad53 at (S/T)Q residues upon replication stress. Phosphorylated Mrc1 functions to stabilize Mec1 presence at the perturbed fork, support Mec1-mediated activation of Rad53, and antagonizes stimulation of the unwinding activity of the Cdc45-MCM-GINS (CMG) complex by Mrc1 (Alcasabas *et al*, 2001; McClure & Diffley, 2021; Naylor *et al*, 2009; Osborn & Elledge, 2003). In addition, Mrc1 functions in the fork protection complex to stabilize stalled replication forks during replication stress (Katou *et al*, 2003; Lou *et al*, 2008; Tourriere *et al*, 2005). Apart from Mrc1, the fork protection complex consists of the Csm3 and Tof1 proteins, both of which are required for Mrc1 presence at the replication fork (Bando *et al*, 2009; Uzunova *et al*, 2014). Finally, Mrc1 is required at the replication fork to promote efficient DNA synthesis during unperturbed replication (Gispan *et al*, 2014; Hodgson *et al*, 2007; Szyjka *et al*, 2005; Yeeles *et al*, 2017).

To resume DNA synthesis after replication fork stalling, the replication checkpoint must be deactivated. At least two parallel mechanisms exist to promote replication checkpoint recovery. One is the de-phosphorylation of Rad53 catalysed by the Pph3- Psy2 phosphatase (O’Neill *et al*, 2007). The other is the proteasome-mediated degradation of Mrc1 (Chaudhury & Koepp, 2017; Fong *et al*, 2013). Degradation of Mrc1 depends in part on ubiquitylation catalysed by the replication fork-associated ubiquitin ligase Skp1-Cullin-F-Box (SCF)-Dia2, which is itself stabilized upon activation of the replication checkpoint (Fong *et al*., 2013; Kile & Koepp, 2010; Mimura *et al*, 2009; Morohashi *et al*, 2009).

We have previously shown that Mrc1 localizes to the intranuclear quality control compartment (INQ) upon methyl methanesulfonate (MMS)-induced replication stress and proteasomal inhibition (Gallina *et al*, 2015). INQ is a perinuclear structure that forms in response to different cellular stresses. Upon MMS-induced replication stress, INQ contains misfolded proteins, proteasomes and 22 additional proteins of various cellular functions, including Pph3 and the nuclear chaperone Btn2. Along with another chaperone named Hsp42, Btn2 is required for Mrc1 localization to INQ as well as for directing misfolded and other INQ substrates to the inclusion (Gallina *et al*., 2015; Miller *et al*, 2015). Interestingly, Mrc1 re-localization to INQ upon MMS-induced replication stress also depends on Dia2 (Gallina *et al*., 2015). It is possible that INQ similarly to other membrane-less structures is governed by liquid-liquid phase separation (Brangwynne *et al*, 2015).

Cdc48 is a conserved homohexameric, ring-shaped AAA+ (extended family of ATPase associated with various activities) ATPase, which functions as a chaperone in various cellular processes, including endoplasmatic reticulum-associated degradation (ERAD), membrane fusion, autophagy, cell cycle regulation and replication stress (Baek *et al*, 2013; Jentsch & Rumpf, 2007; Ramadan *et al*, 2017). The intrinsic ATPase activity of Cdc48 is required for many of these functions but substrate recognition and processing are achieved mostly by Cdc48-associated cofactors (Baek *et al*., 2013; Hartmann-Petersen *et al*, 2004; Jentsch & Rumpf, 2007; Meyer *et al*, 2002). Targeting of ubiquitylated substrates by Cdc48 and its associated cofactors mainly results in delivery of the substrate to the proteasome for degradation (Baek *et al*., 2013; Baek *et al*, 2011; Elsasser & Finley, 2005; Verma *et al*, 2011). However, Cdc48 has also been shown to mediate chromatin extraction and disassembly of ubiquitylated protein complexes (Maric *et al*, 2014; Ramadan *et al*., 2017; Rape *et al*, 2001). For example, Cdc48 promotes chromatin eviction of cohesin at stalled replication forks (Frattini *et al*, 2017) and disassembly of CMG upon replication termination (Maric *et al*., 2014; Maric *et al*, 2017; Moreno *et al*, 2014).

In this study, we show that Btn2 and Cdc48 are instrumental for replication restart following MMS-induced replication stress. Mrc1 is recognized by the Cdc48 cofactors Ufd1 and Otu1 and jointly, Cdc48-Ufd1 and -Otu1 function to clear Mrc1 from INQ upon recovery from MMS-induced replication stress. Mrc1 interacts with the INQ chaperone Btn2 upon MMS treatment and ectopic expression of Btn2 is sufficient to sequester Mrc1 in INQ. Importantly, in the absence of Cdc48, Btn2-mediated sequestering of Mrc1 in INQ supports replication recovery. Our findings point to INQ as a hub for Cdc48-mediated regulation of Mrc1 during replication stress.

## Results

### Mrc1 is transiently sequestered in INQ during replication stress

We have previously shown that Mrc1 and 22 other proteins localize to the perinuclear structure INQ upon MMS-induced replication stress (Gallina *et al*., 2015) and decided to analyse the functional significance of this re-localization in relation to the replication checkpoint, of which Mrc1 is the primary mediator (Alcasabas *et al*., 2001; Osborn & Elledge, 2003). To this end, localization of Mrc1 was observed in cells expressing a C-terminally yellow fluorescent protein (YFP)-tagged Mrc1 protein (Gallina *et al*., 2015). Mrc1-YFP was visualized in cells synchronized in G1 by addition of alpha factor, during release into S phase with or without MMS-induced replication stress and during recovery after removal of MMS (Figure 1A). Upon release from alpha factor arrest into MMS, Mrc1-YFP re-localized from a pan-nuclear pattern to a single perinuclear focus in a subset (10%) of cells (Figure 1A). These foci on average contain 20% of nuclear Mrc1 (Figure 1B). When cells were subsequently allowed to recover from MMS-induced replication stress, Mrc1-YFP redistributed back into a pan-nuclear pattern within 60 minutes.

**Figure 1.**
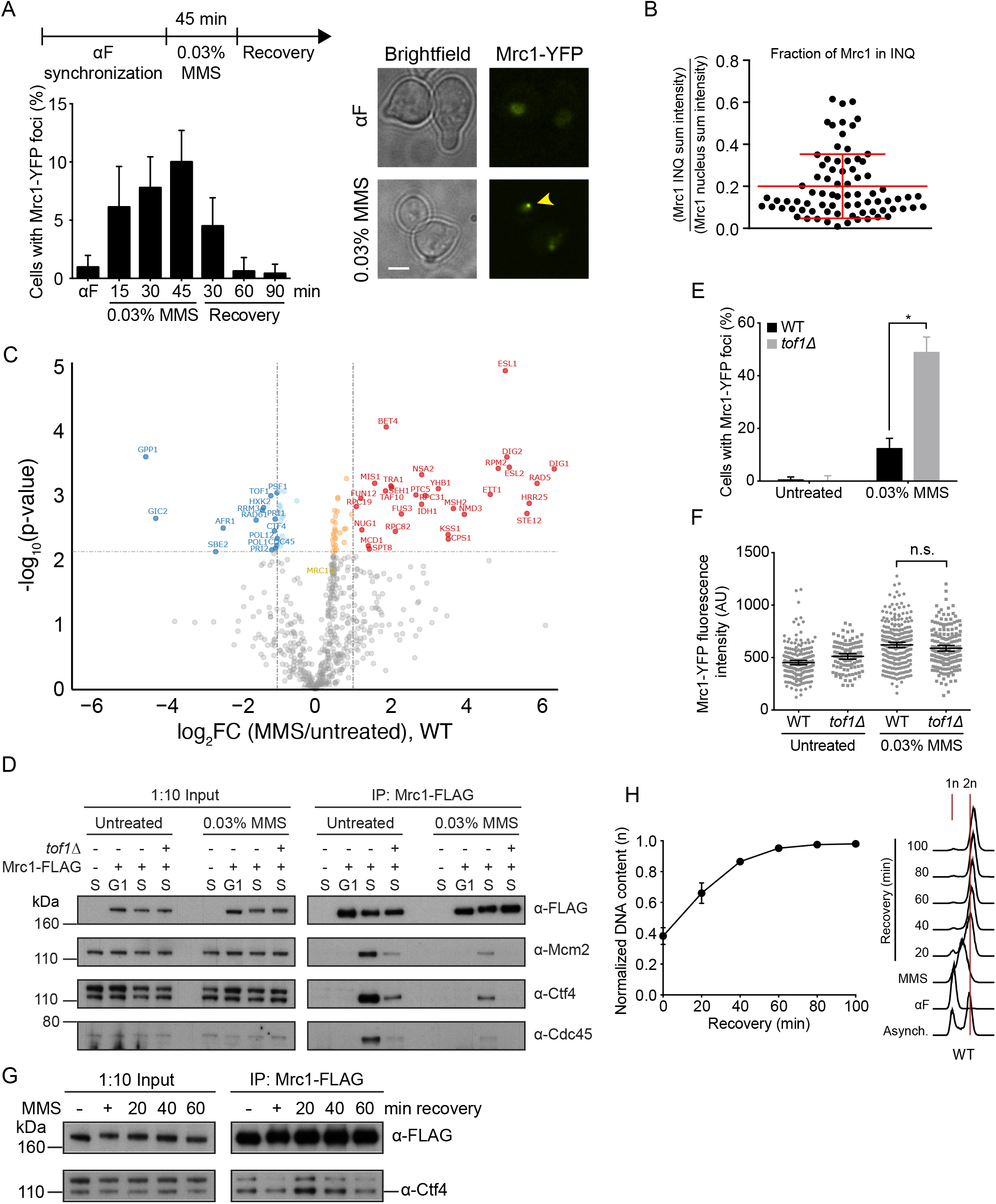
Mrc1 localization to and disappearance from INQ correlates with Mrc1 dissociation from the fork. **A.** Temporal dissection of Mrc1 localization to INQ. *Top left*, schematic representation of the workflow used in this study. Unless otherwise stated, cells were grown to exponential phase and synchronized by addition of alpha factor (αF) before being released for 45 minutes into 0.03% MMS. Finally, cells were allowed to recover in medium without MMS. *Bottom left*, graph showing the percentage of WT cells expressing Mrc1-YFP (IG147) that contain foci at the indicated time points. Two replicates and minimum 250 cells were analysed for each time point. Error bars represent 95% confidence intervals. *Right*, representative images of Mrc1-YFP localization after alpha factor arrest and 45 minutes after release from alpha factor into 0.03% MMS. Arrowhead points to Mrc1-YFP INQ focus. Scale bar, 3 µm. **B.** Quantitation of Mrc1 in foci. Cells expressing Mrc1-YFP (IG147) were released from alpha factor synchronization into 0.03% MMS and imaged after 45 minutes of incubation. The images were used to quantify both the INQ and the nuclear signal and the fraction of total nuclear YFP signal present in the INQ foci was calculated. The error bars represent the mean and standard deviation. 78 cells were measured. **C.** Proteomics analysis of Mrc1 interacting proteins during replication stress. Wild-type cells (ML1168-1B) were arrested in G1 phase using alpha-factor pheromone and released into S phase with or without 0.03% MMS. Cells were harvested for protein extraction 25 min (without MMS) or 45 min (with MMS) after release from alpha-factor arrest, and after 30 min of recovery in medium without MMS. After immunoprecipitation of Mrc1, co-immunoprecipitating proteins were identified by LC-MS/MS. Proteins that display a significantly reduced or increased interaction with Mrc1 in MMS relative to untreated (left) are highlighted in blue and red, respectively. Proteins that display a significantly increased interaction with Mrc1 during recovery relative to MMS (right) are highlighted in red. Dashed lines indicate thresholds (-1 > log2FC (MMS/untreated) > 1; -log10(p-value) for the protein with the highest significant posthoc adjusted p-value below 0.05). Proteins with significant adjusted p-values but -1 < log2FC (MMS/untreated) < 1 are colored in light blue and orange, respectively. Mrc1 is highlighted in yellow. **D.** Mrc1 interacts with several replication fork components in S phase. Input and Mrc1-FLAG immunoprecipitation samples were prepared from WT cells expressing Mrc1-FLAG (CC201-7B) synchronized by alpha factor (G1), released from alpha factor into fresh medium for 20 minutes or into medium containing 0.03% MMS for 45 minutes (S). In addition, samples were prepared from untreated S-phase WT cells (CC202-1A) and from *tof1*Δ cells expressing Mrc1-FLAG (CC215-41B) released from alpha factor arrest for 20 minutes into fresh medium and for 45 minutes into 0.03% MMS. Mrc1-FLAG and the replication fork components Mcm2, Ctf4 and Cdc45 were analysed by immunoblotting. **E.** Mrc1 localization to INQ is enhanced in the absence of *TOF1*. Asynchronous WT or *tof1Δ* cells (IG147 and CC133-1A) were imaged untreated or after 60 minutes treatment with 0.03% MMS. Percentages are averages of two experiments and minimum 150 cells were counted. Error bars represent 95% confidence intervals. *, P < 0.0001. **F**. *TOF1* does not affect nuclear Mrc1 protein level during replication stress. The fluorescence intensity of the Mrc1- YFP nuclear signal was measured from images of the experiment described in E. The line across the scatter plot represents the mean and the error bars represent 95% confidence intervals. A minimum of 100 cells was measured. **G.** Temporal dissection of Mrc1 interaction with the replisome. Input and Mrc1-FLAG immunoprecipitation samples were prepared from WT cells (CC201-7B) harvested 20 minutes after release from alpha factor arrest in the absence of MMS (-) or 45 minutes after release into 0.03% MMS (+) and 20, 40 and 60 minutes into recovery from MMS. Mrc1-FLAG and the replication fork component Ctf4 were analysed by immunoblotting. **H.** Replication recovery correlates with Mrc1 disappearance from INQ. WT cells (CC44-8A) were harvested before and after alpha factor arrest, after 45 minutes release into 0.03% MMS and after 20, 40, 60, 80 and 100 minutes recovery from MMS in medium containing nocodazole. *Top*, graph showing the normalized DNA content as measured from the flow cytometry data in samples taken immediately before (t = 0, MMS) and at indicated time points after recovery from MMS. Values are averages of 8 experiments and error bars represent standard error of the mean (SEM). *Bottom*, representative propidium iodide (PI) profiles. Red lines indicate peaks of 1n and 2n DNA content.

As Mrc1 is a replication fork associated protein (Katou *et al*., 2003; Lou *et al*., 2008; Osborn & Elledge, 2003), we hypothesized that the interaction between Mrc1 and replisome components would change upon localization of Mrc1 to INQ. To examine the interactome of Mrc1 in an unbiased manner, we performed immunoprecipitation of Mrc1-YFP followed by identification of co-immunoprecipitated proteins by LC-MS/MS in an unchallenged S phase (Reusswig *et al*, 2021), or in an MMS challenged S phase and upon recovery from MMS treatment. This approach confirmed that Mrc1 interacts with a number of replisome components in an unchallenged S phase (Table S6) and revealed a reduced interaction with replication factors in MMS such as Cdc45, MCM (Mcm2-7), GINS (Sld5, Psf1, Psf2, Psf3), Csm3, Tof1, Ctf4, polymerase alpha (Pri1, Pri2, Pol12, and Pol1), and polymerase epsilon (Pol2, Dpb2, Dpb3) (Figure 1C, Figure S1, Figure S2 and Table S7). These interactions were partially regained during recovery from MMS (Figure S1). The transient dissociation of replisome components Mcm2, Ctf4 and Cdc45 from Mrc1 was confirmed by Western blotting of immunoprecipitated C-terminally FLAG-tagged Mrc1 (Figure 1D). Moreover, in an MMS-stressed S phase, Mrc1-FLAG interaction with the fork components was reduced to the same degree as in an untreated *tof1Δ* mutant, in which Mrc1 localization to the fork has been shown to be compromised (Bando *et al*., 2009; Uzunova *et al*., 2014), and completely abolished in the MMS-treated *tof1*Δ mutant. Concurrently, formation of Mrc1 foci rose to a four-fold higher level in MMS-treated *tof1*Δ mutant cells compared to that observed in wild-type cells (Figure 1E). This was not due to a change in the Mrc1 protein level (Figure 1F). During recovery, Mrc1 exhibited increased interaction with the 26S proteasome subunits Rpn1, Rpn3, Rpn8 and Rpn11 (Figure S1 and Table S6), which is consistent with proteasomal degradation of Mrc1 in response to replication stress (Chaudhury & Koepp, 2017). Finally, upon recovery from MMS, Mrc1-FLAG interaction with Ctf4 was restored and subsequently decreased with time (Figure 1G), presumably as DNA replication gradually completed. Thus, in wild-type cells, localization of Mrc1 to INQ is inversely correlated with its interaction with the replisome during MMS treatment and during recovery from MMS-induced replication stress.

Next, we analyzed the temporal localization of Mrc1 to INQ relative to completion of replication after MMS treatment by flow cytometry of propidium iodide (PI)-stained cells. Cells were allowed to recover from MMS in medium containing nocodazole to prevent cells from dividing and commencing a new cell cycle. When recovering from an MMS-stressed S phase, the data showed a gradual increase in the fraction of cells with a 2n DNA content (Figure 1H), representing cells in which replication had completed. 60 minutes into recovery, 85% of cells had completed DNA replication. Taken together, these data show that Mrc1 disappearance from INQ and reappearance at the replication fork correlates with resumption of replication.

### The Btn2 chaperone sequesters Mrc1 in INQ

We have previously shown that the INQ-localized chaperone Btn2 is required for Mrc1 re-localization to INQ (Gallina *et al*., 2015). Therefore, in order to test for a physical connection between Mrc1 and Btn2, co-immunoprecipitation of Btn2-HA with Mrc1- FLAG was performed. While no interaction was detected during an unperturbed S-phase, MMS treatment led to both an increase in Btn2 protein level and interaction with Mrc1 (Figure 2A). In order to visualize this interaction *in situ*, we took advantage of the bimolecular fluorescence complementation (BiFC) assay, in which N-terminal (VN) and C-terminal (VC) non-fluorescent fragments of the Venus fluorescent protein were fused to Btn2 and Mrc1, respectively, and their physical association evaluated by the appearance of a fluorescence signal (Figure 2B). In agreement with the co-immunoprecipitation data, no BiFC signal was detected in untreated cells, while a perinuclear focal fluorescence signal colocalizing with the INQ marker Cmr1 was detected in a subset of cells upon MMS treatment (Figure 2C and 2D). The subset of cells containing this INQ-localized Mrc1-Btn2 interaction was drastically increased upon additional treatment with the proteasome inhibitor MG132 (Figure 2C and 2D).

**Figure 2.**
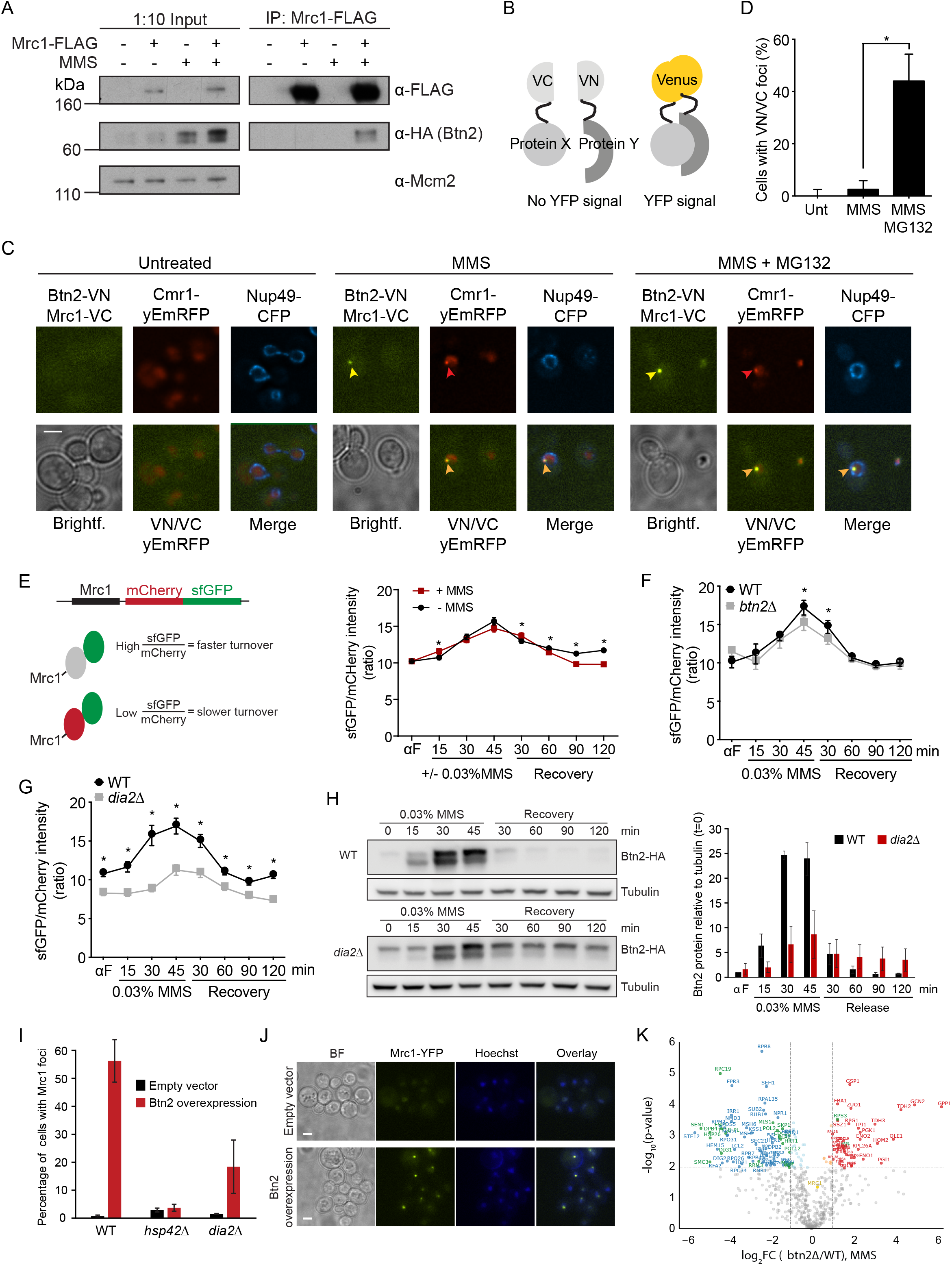
Btn2 interacts with and targets Mrc1 to INQ and facilitates Mrc1 turnover. **A.** Btn2 interacts with Mrc1 during MMS-induced replication stress. Input and Mrc1-FLAG immunoprecipitation samples were prepared from cells expressing Btn2-HA with or without Mrc1-FLAG (CC201-11A and CC223-35B) harvested 20 minutes after release from alpha factor arrest in the absence of MMS (-) or 45 minutes after release into 0.03% MMS (+). Btn2-HA and Mrc1-FLAG were analysed by immunoblotting. Mcm2 was used as a loading control for input samples. **B.** Schematic representation of the principle of bimolecular fluorescence complementation (BiFC). Interaction of VC- and VN-tagged proteins X and Y results in the reconstitution of fluorescent Venus protein. **C-D.** Mrc1 and Btn2 interact in INQ after replication stress. Cells expressing Mrc1-VC, Btn2-VN and Cmr1-yEmRFP (CC299-12C) were transformed with pML4 for expression of Nup49-CFP and imaged untreated and after 2 hours 0.05% MMS treatment with or without MG132. **C**. Representative images from the experiment. Arrowheads point to Mrc1/Btn2 interactions in INQ foci. Scale bar, 3 µm. **D**. Quantitation of the experiment performed in **C**. A minimum of 100 cells was analysed for each treatment. Error bars represent 95% confidence intervals. *, P < 0.0001. **E.** Mrc1 turnover correlates with localization in INQ. *Left*, schematic representation of the Mrc1-mCherry-sfGFP timer construct. *Right*, turnover of the Mrc1 timer construct in WT cells (CC98) was analysed as the ratio between the fluorescence intensity of the two fluorophores as measured from images acquired at the time points indicated. A representative experiment out of four is shown. A minimum of 75 cells were measured for each time point. Error bars represent 95% confidence intervals. **F.** *BTN2* facilitates Mrc1 turnover after replication stress. The ratio of the fluorescence intensity of mCherry and sfGFP was measured in images of WT and *btn2Δ* cells expressing Mrc1-mCherry-sfGFP (CC98 and CC102-9C) acquired at the time points indicated. A representative experiment out of two is shown. A minimum of 50 cells was measured for each time point and error bars represent 95% confidence intervals. *, P < 0.01. **G.** Mrc1 turnover is impaired in the absence of *DIA2*. The ratio of the fluorescence intensity of mCherry and sfGFP was measured in images of WT and *dia2Δ* cells expressing Mrc1-mCherry-sfGFP (CC98 and CC101-8C) acquired at the time points indicated. A representative experiment out of two is shown. A minimum of 75 cells were measured for each time point and error bars represent 95% confidence intervals. *, P < 0.0001. **H.** Dia2 promotes accumulation of Btn2. *Left,* representative immunoblot of whole cell extracts prepared from WT (CC201-7D) and *dia2Δ* (JA29- 3B) cells with HA-tagged Btn2, harvested at the indicated time points after release from alpha factor arrest into 0.03% MMS and during recovery from MMS treatment. Btn2 protein level was analyzed by immunoblotting with tubulin used as loading control. *Right*, quantitation of the data in the left panel with the values representing tubulin- normalized Btn2-HA protein levels relative to alpha factor arrested cells. Shown is the mean of two experiments. Error bars represent the standard deviation. **I.** Ectopic Btn2 expression sequesters Mrc1 in INQ in the absence of DNA damage. WT (IG147), *hsp42Δ* (CC2-6B), and *dia2Δ* (CC37-5A) cells were transformed with either empty vector (pRS425) or a vector (pRS425-Btn2) for galactose inducible Btn2 expression. Images of the transformed cells after a 1-hour incubation with 2% galactose were used to quantify the percentage of cells with Mrc1-YFP foci. Shown is the mean of three experiments. The error bars represent the standard deviation. **J.** Btn2 sequesters Mrc1 in INQ. Representative images of the experiment in panel **I**. Scale bar, 3 µm. **K.** Proteomics analysis of Mrc1 interacting proteins in replication stressed *btn2*Δ cells. WT and *btn2*Δ cells (ML1168-1B and ML1167-10D) were arrested in G1 phase using alpha-factor and released into S phase with 0.03% MMS. Cells were harvested for protein extraction 45 min after release from alpha-factor arrest into MMS. After immunoprecipitation of Mrc1, co-immunoprecipitating proteins were identified by LC- MS/MS. Proteins that display a significantly reduced interaction with Mrc1 in MMS- treated *btn2*Δ cells relative to MMS-treated wild-type cells are highlighted in green and blue, depending on whether or not the protein was already identified as a significant interactor of Mrc1 in wild-type cells under the same condition (Table S6), respectively. Proteins that display a significantly increased interaction with Mrc1 in MMS-treated *btn2*Δ cells relative to MMS-treated wild-type cells are highlighted in green and red, depending on whether or not the protein was already identified as a significant interactor of Mrc1 in wild-type cells under the same condition (Table S6), respectively. Dashed lines indicate thresholds (-1 > log2FC (*btn2*Δ/WT) > 1; -log10(p-value) for the protein with the highest significant posthoc adjusted p-value below 0.05). Proteins with significant adjusted p-values but -1 < log2FC (*btn2*Δ/WT) < 1 are colored in light blue and orange, respectively. Mrc1 is highlighted in yellow.

In light of these data, as well as our previous findings that a subset of INQ structures contain proteasomes and that Mrc1 accumulates in INQ upon proteasomal inhibition (Gallina *et al*., 2015), we hypothesized that replication stress-induced Mrc1 localization to and interaction with Btn2 in INQ are coupled to proteasomal degradation of Mrc1. To investigate this, Mrc1 turnover during and upon recovery from MMS-induced replication stress was analysed using a fluorophore-based timer construct, in which Mrc1 is tagged in tandem by the slow-maturing fluorophore mCherry and the fast-maturing fluorophore sfGFP. The ratio between the fluorescence intensities of the two fluorophores can be used to compare the age of the tagged protein under different conditions (Khmelinskii *et al*, 2012). Mrc1-mCherry-sfGFP turnover increased upon release from alpha factor arrest into an MMS-perturbed S phase and subsequently decreased back to the base line level within 60 minutes of recovery from MMS (Figure 2E), correlating with the temporal profile of Mrc1 localization in INQ. However, the Mrc1 turnover profiles were similar in the presence and absence of MMS, suggesting that Mrc1 turnover does not increase further upon MMS treatment. More likely, the increase sfGFP/mCherry ratio observed in S phase is due to increased *de novo* expression of Mrc1 in S phase after mRNA levels have peaked in late G1 (Cho *et al*, 1998; de Lichtenberg *et al*, 2003; Pramila *et al*, 2002; Spellman *et al*, 1998). In parallel, the total Mrc1 protein level was analysed by immunoblotting of whole cell extracts in cells expressing a FLAG-tagged Mrc1 protein from the endogenous locus. During MMS-induced replication stress, Mrc1-FLAG protein level remained constant relative to tubulin with a small transient decrease during recovery from MMS (Figure S3A). This is consistent with the report that MMS treatment does not induce *MRC1* gene expression (Gasch *et al*, 2001; Pramila *et al*, 2006). Therefore, we conclude that increased degradation of Mrc1 protein in S phase is balanced by *de novo* synthesis. In the *btn2Δ* mutant, turnover of Mrc1-mCherry-sfGFP was mildly decreased compared to WT (Figure 2F), indicating that degradation of Mrc1 is only mildly stimulated by Btn2. By contrast, deletion of *DIA2*, which is an SCF ubiquitin ligase-associated F-box protein that localizes to the replication fork and targets Mrc1 for ubiquitylation and proteasomal degradation during unperturbed replication and during recovery from MMS-induced replication stress (Fong *et al*., 2013; Mimura *et al*., 2009; Morohashi *et al*., 2009), dramatically reduced the turnover of Mrc1 (Figure 2G), suggesting that Dia2 promotes an intrinsic destabilization of Mrc1 even in the absence of MMS treatment (Fong *et al*., 2013; Mimura *et al*., 2009). Since Dia2 has also been reported to facilitate inducible gene expression (Andress *et al*, 2011), we considered the possibility that MMS-induced accumulation of Btn2 is dependent on *DIA2.* Indeed, Btn2 accumulation is deregulated in *dia2*Δ cells with a higher constitutive level but less robust accumulation upon addition of MMS (Figure 2H). This may contribute to the reduction in Mrc1 foci observed in *dia2*Δ cells (Gallina *et al*., 2015).

To test if expression of Btn2 is sufficient to sequester Mrc1 in INQ in the absence of DNA damage, we ectopically expressed Btn2 from a galactose-inducible promoter in WT, *hsp42*Δ and *dia2*Δ cells (Figure 2I-J). This experiment showed that merely overexpressing Btn2 results in the re-localization of Mrc1 to INQ, and this re- localization depends on *HSP42* and *DIA2* to the same extent as the MMS-induced re- localization of Mrc1 to INQ (Gallina *et al*., 2015), suggesting that Mrc1 ubiquitination by Dia2 is still required for re-localization of Mrc1 to INQ, when Btn2 is overexpressed. Consistently, Hsp42 and Btn2 interact in the BiFC assay (Figure S3B). Similarly, Btn2 overexpression also induced formation of Pph3 and Cmr1 foci, which have previously been shown to re-localize to INQ after MMS-treatment or inhibition of the proteasome (Gallina *et al*., 2015) (Figure S3C). While both Pph3 and Cmr1 required *HSP42* for re- localization to foci upon Btn2 overexpression, only Pph3 required *DIA2* (Figure S3C). This may reflect a requirement for ubiquitylation by SCF-Dia2 for the recruitment of some proteins to INQ. Consistent with previous studies (Chaudhury & Koepp, 2017), deletion of *DIA2* resulted in a significant delay in replication completion upon recovery from MMS (Figure S3D) as well as an overall slower intrinsic replication during an unperturbed S phase (Figure S3E). Collectively, these findings suggest that Mrc1 interacts directly with Btn2, and Btn2 expression promotes the sequestration of Mrc1 in INQ even in the absence of exogenous DNA damage.

To determine how the interactome of Mrc1 changes in *btn2*Δ cells, which cannot recruit Mrc1 to INQ during replication stress, we conducted proteomics analysis as described above using a *btn2*Δ mutant strain expressing Mrc1-YFP. This analysis revealed that in all conditions, Mrc1 had reduced interaction with replication factors (Figure 2K, S1 and S3F, and Table S7). Furthermore, the interaction of Mrc1 to the SCF-Dia2 ubiquitin ligase complex (Skp1-Cdc53-Dia2-Hrt1) was reduced in MMS and during recovery from MMS in the *btn2*Δ mutant (Figure S2C), suggesting that localization of Mrc1 to INQ enhances its targeting for degradation.

### Cdc48 is required for Mrc1 turnover and replication recovery

To further dissect the mechanism responsible for Mrc1 re-localization between the replication fork and INQ during MMS-induced replication stress, we investigated the potential role of another chaperone, Cdc48. Cdc48 has been associated with the extraction of proteins from chromatin in several studies (reviewed in (Ramadan *et al*., 2017)). Of special interest was the recent work, showing that Mcm7 of the CMG helicase is ubiquitylated by SCF-Dia2 upon replication termination and that the CMG helicase is subsequently recognized and extracted from chromatin by Cdc48-Ufd1- Npl4 (Maculins *et al*, 2015; Maric *et al*., 2014; Maric *et al*., 2017). We hypothesized that Cdc48 could play a similar role in extraction of SCF-Dia2-ubiquitylated Mrc1 from the replication fork during replication stress. As *CDC48* is an essential gene, we took advantage of the auxin-induced degradation system (Nishimura *et al*, 2009) to conditionally eliminate the Cdc48 protein. In order to investigate whether Cdc48 is required for re-localizing Mrc1 from the replication fork to INQ upon MMS-induced replication stress, the localization of Mrc1-YFP to INQ was examined in WT and cells expressing Cdc48-aid with or without auxin in untreated and MMS-treated conditions. To our surprise, there was no defect in Mrc1-YFP localization to INQ in the absence of Cdc48 (Figure 3A). On the contrary, we observed a marked increase in the percentage and intensity of Mrc1-YFP foci as well as a persistence of foci up to 2 hours into recovery from MMS. This indicates that Cdc48 is not involved in extracting Mrc1 from the replication fork, when Mrc1 re-localizes to INQ, but rather that Cdc48 could be required for clearing Mrc1 from INQ after MMS-induced replication stress. In fact, Cdc48 may have a more general function in extracting proteins from INQ, as spontaneous and MMS-induced foci of Cmr1-YFP and Pph3-YFP, two other replication checkpoint associated proteins that localize to INQ (Gallina *et al*., 2015), also persisted in INQ upon auxin-induced degradation of Cdc48 (Figure S4A).

**Figure 3.**
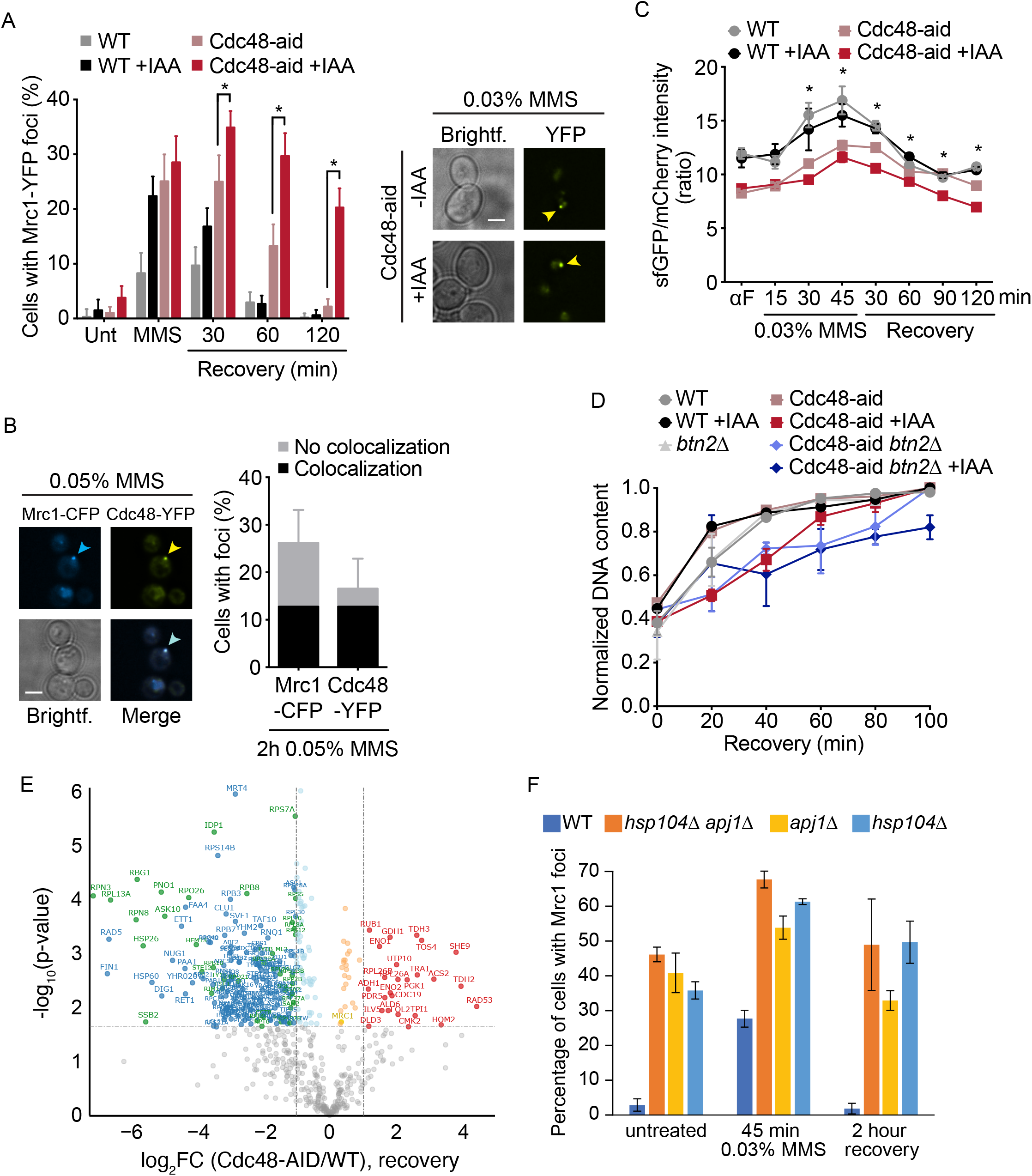
Cdc48 is required for Mrc1 turnover and replication recovery. **A.** Mrc1 persists in INQ in the absence of Cdc48. *Left*, graph showing the percentage of asynchronous WT and Cdc48-aid cells (IG147 and CC127-4A) with or without auxin (Indole-3-acetic acid, IAA) displaying Mrc1-YFP foci when untreated (Unt), treated with 0.03% MMS for 45 minutes and after 30, 60, and 120 minutes recovery from MMS. Values are averages of two experiments. Minimum 300 cells were analysed and error bars represent 95% confidence intervals. *, P < 0.001. *Right*, Representative images of Mrc1-YFP foci in Cdc48-aid cells treated for 45 minutes with 0.03% MMS with or without auxin (IAA). Arrowheads point to Mrc1-YFP INQ foci. Scale bar, 3 µm. **B.** Cdc48 and Mrc1 colocalize in asynchronous cells after MMS-induced replication stress. *Left*, representative image of Mrc1-CFP and Cdc48-YFP colocalization after 2 hours of 0.05% MMS treatment (CC182-10B). Arrowheads point to Cdc48/Mrc1 colocalization foci. Scale bar, 3 µm. *Right*, quantitation of Cdc48/Mrc1 colocalization foci. Values are averages of two experiments and 290 cells were analysed. Error bars represent 95% confidence intervals. **C.** Mrc1 turnover is impaired in the absence of Cdc48. The ratio of the fluorescence intensity of mCherry and sfGFP was measured in WT and Cdc48-aid cells expressing Mrc1-mCherry-sfGFP (CC98 and CC132-13C) with and without auxin at the time points indicated. A representative experiment out of two is shown. A minimum of 75 cells were measured and error bars represent 95% confidence intervals. *, P < 0.0001. **D.** Replication recovery after MMS-induced replication stress is impaired in the absence of Cdc48 and is synthetically compromised in the absence of Cdc48 and *BTN2.* WT, Cdc48-aid (CC44-8A and CC128-33A) with or without auxin, Cdc48-aid *bnt2Δ* (CC136-29D) with or without auxin and *btn2Δ* cells (CC26-11B) were harvested before and after alpha factor arrest, after 45 minutes release into 0.03% MMS and after 20, 40, 60, 80, and 100 minutes recovery from MMS in medium containing nocodazole. The graphs show the normalized DNA content as measured from the flow cytometry data in samples taken immediately before and at indicated time points during recovery from MMS. Values are averages of two to 8 experiments and error bars represent SEM. **E.** Proteomics analysis of Mrc1 interacting proteins in replication-stressed Cdc48-depleted cells. WT and *CDC48-AID* cells (ML1168-1B and ML1166-33A) were arrested in G1 phase using alpha-factor pheromone, Cdc48 was depleted by addition of auxin in parallel with the alpha factor, and released into S phase with 0.03% MMS for 45 min. Cells were harvested for protein extraction after 30 min of recovery in medium without MMS. After immunoprecipitation of Mrc1, co-immunoprecipitating proteins were identified by LC-MS/MS. Proteins that display a significantly reduced interaction with Mrc1 during recovery of Cdc48-depleted cells relative to wild-type cells are highlighted in green and blue, depending on whether or not the protein was already identified as a significant interactor of Mrc1 in wild-type cells under the same condition (Table S6), respectively. Proteins that display a significantly increased interaction with Mrc1 during recovery of Cdc48-depleted cells relative to wild-type cells are highlighted in green and red, depending on whether or not the protein was already identified as a significant interactor of Mrc1 in wild-type cells under the same condition (Table S6), respectively. Dashed lines indicate thresholds (-1 > log2FC (Cdc48-AID/WT) > 1; -log10(p-value) for the protein with the highest significant posthoc adjusted p-value below 0.05). Proteins with significant adjusted p-values but -1 < log2FC (MMS/untreated) < 1 are colored in light blue and orange, respectively. Mrc1 is highlighted in yellow. **F.** INQ-localized Mrc1 is a substrate for Apj1 and Hsp104 chaperones. Wild-type (IG147), *hsp104*Δ (ML1172- 11C), *apj1*Δ (ML1172-6D) and *hsp104Δ apj1*Δ (ML1172-1C) cells were grown to exponential phase in SC+Ade at 25°C and imaged by microscopy untreated, after 45 min incubation with 0.03% MMS, and after 2 hours of recovery from MMS (N = 2-3). A minimum of 290 cells were analysed per strain. Error bars represent the standard deviation.

Cdc48 has been shown to interact with and deliver ubiquitylated proteins to the proteasome for degradation (Baek *et al*., 2011; Dai *et al*, 1998; Verma *et al*, 2000; Verma *et al*., 2011). As the proteasome localizes to INQ in a subset of cells (Gallina *et al*., 2015) and Mrc1 is degraded during S-phase checkpoint recovery (Figure S3A) (Fong *et al*., 2013), we hypothesized that Cdc48 may recognize ubiquitylated Mrc1 in INQ and deliver the protein for proteasomal degradation here, thus explaining the persistence of Mrc1 foci in the absence of Cdc48. To test this hypothesis, we first investigated whether Cdc48 itself is present in INQ. When imaging cells expressing Mrc1-CFP and Cdc48-YFP after 2 hours of MMS treatment, 77% of the observed Cdc48-YFP foci colocalized with Mrc1-CFP while 49% of the observed Mrc1-CFP foci colocalized with Cdc48-YFP (Figure 3B). Consistently, our proteomics analysis consistently showed co-immunoprecipitation of Mrc1 and Cdc48, which was abolished upon auxin-induced depletion of Cdc48 (Figure S4D). We therefore pursued the hypothesis that Cdc48 targets Mrc1 for proteasomal degradation following MMS-induced replication stress by analysing Mrc1-mCherry-sfGFP turnover during and in recovery from MMS treatment in WT and cells expressing Cdc48-aid with or without auxin. Auxin alone had little effect on the temporal profile of Mrc1-mCherry-sfGFP turnover in WT cells, while Cdc48-aid significantly decreased turnover during the entire time course and addition of auxin to Cdc48-aid cells further reduced Mrc1 turnover (Figure 3C). This indicates that functional Cdc48 is instrumental for degradation of Mrc1.

To investigate the relevance of Cdc48 for recovery from the replication checkpoint, the kinetics of replication completion was examined in the presence or absence of Cdc48 using flow cytometry. Auxin-induced degradation of Cdc48-aid delayed replication recovery with only 67% of these cells having reached a 2n DNA content after 40 minutes of release from MMS treatment compared to 89% for WT cells (Figure 3D and S4C). This was not due to intrinsically slower replication in the absence of Cdc48, as auxin-treated cells expressing Cdc48-aid completed replication with the same kinetics as WT when released from alpha factor arrest into an unperturbed S phase (Figure S3E). Furthermore, auxin or expression of Cdc48-aid alone did not compromise replication recovery after MMS-induced stress, as these cells completed replication with the same kinetics as non-auxin-treated WT cells (Figures 3D and S4C). Additionally, while having no significant effect on replication completion after MMS-induced replication stress alone, deletion of *BTN2* exacerbated the delay observed in auxin-treated Cdc48-aid cells (Figure 3D and S4C). This suggests that when Cdc48- mediated degradation of Mrc1 is compromised, sequestration of Mrc1 in INQ promotes replication completion during recovery from MMS-induced replication stress. Collectively, these findings indicate that Cdc48 facilitates turnover of INQ-localized Mrc1 potentially to enable replication completion during recovery from MMS-induced replication stress.

To globally examine the Mrc1 interactome in the absence of Cdc48, we conducted proteomics analysis as described above after depletion of Cdc48. Most notably, we observed a dramatic increase in the interaction of Mrc1 with the Rad53 checkpoint kinase in Cdc48-depleted cells (Figure 3E and S4D), suggesting that checkpoint signaling is increased when Cdc48 fails to release Mrc1 from INQ.

### Disaggregation by Apj1 and Hsp104 chaperones is distinct from Cdc48

A recent study identified Mrc1 as a potential substrate for disaggregation by the Apj1 and Hsp104 chaperones (den Brave *et al*, 2020). Since these chaperones were previously shown to colocalize with Mrc1 in INQ (Gallina *et al*., 2015), we decided to compare the impact of these chaperones to that of Cdc48. First, we examined the localization of Mrc1 to INQ in mutants of the chaperones (Figure 3F). In line with a model where Apj1 and Hsp104 clears substrates from INQ through proteasomal degradation or refolding, respectively (den Brave *et al*., 2020), we observed accumulation of Mrc1 in INQ even in the untreated condition in *apj1*Δ and *hsp104*Δ single and double mutants, which is consistent with Mrc1 exhibiting a high degree of intrinsic disorder (Meszaros *et al*, 2018). Surprisingly, proteomics analysis of proteins co-immunoprecipitating with Mrc1 showed that the interactome of Mrc1 was largely unaffected by mutation of Apj1 and Hsp104 except during recovery, where the interaction of Mrc1 with the proteasome was reduced (Figures S1 and S5E; Table S7). This observation suggest that the functional consequences of Mrc1 aggregation in the absence of Apj1 and Hsp104 might be limited and we therefore decided to focus on the functional implications of Cdc48 and its cofactors on Mrc1 in the remainder of this study.

### Mrc1 interacts with several Cdc48 cofactors

Although Cdc48 can directly bind ubiquitin at the N-terminal domain (Dai & Li, 2001), the contribution of Cdc48 cofactors to recognition of ubiquitylated and SUMOylated substrates appears to be critical for Cdc48 function (Baek *et al*., 2013; Bergink *et al*, 2013; Hartmann-Petersen *et al*., 2004; Meyer *et al*., 2002). In order to identify the Cdc48 cofactors that are important for extracting Mrc1 from INQ, we screened mutants of the 15 known Cdc48 cofactors for persistence of Mrc1 foci 2 hours into recovery from MMS treatment (Baek *et al*., 2013). Gene deletions or chemically induced depletion of the Cdc48 cofactors revealed six disruptions (*otu1Δ*, *ubx3Δ*, *ubx5Δ*, Ufd1- aid*-YFP, *vms1Δ* and *wss1Δ*) that resulted in varying levels of persisting Mrc1 foci (Figure 4A and Table S1).

**Figure 4.**
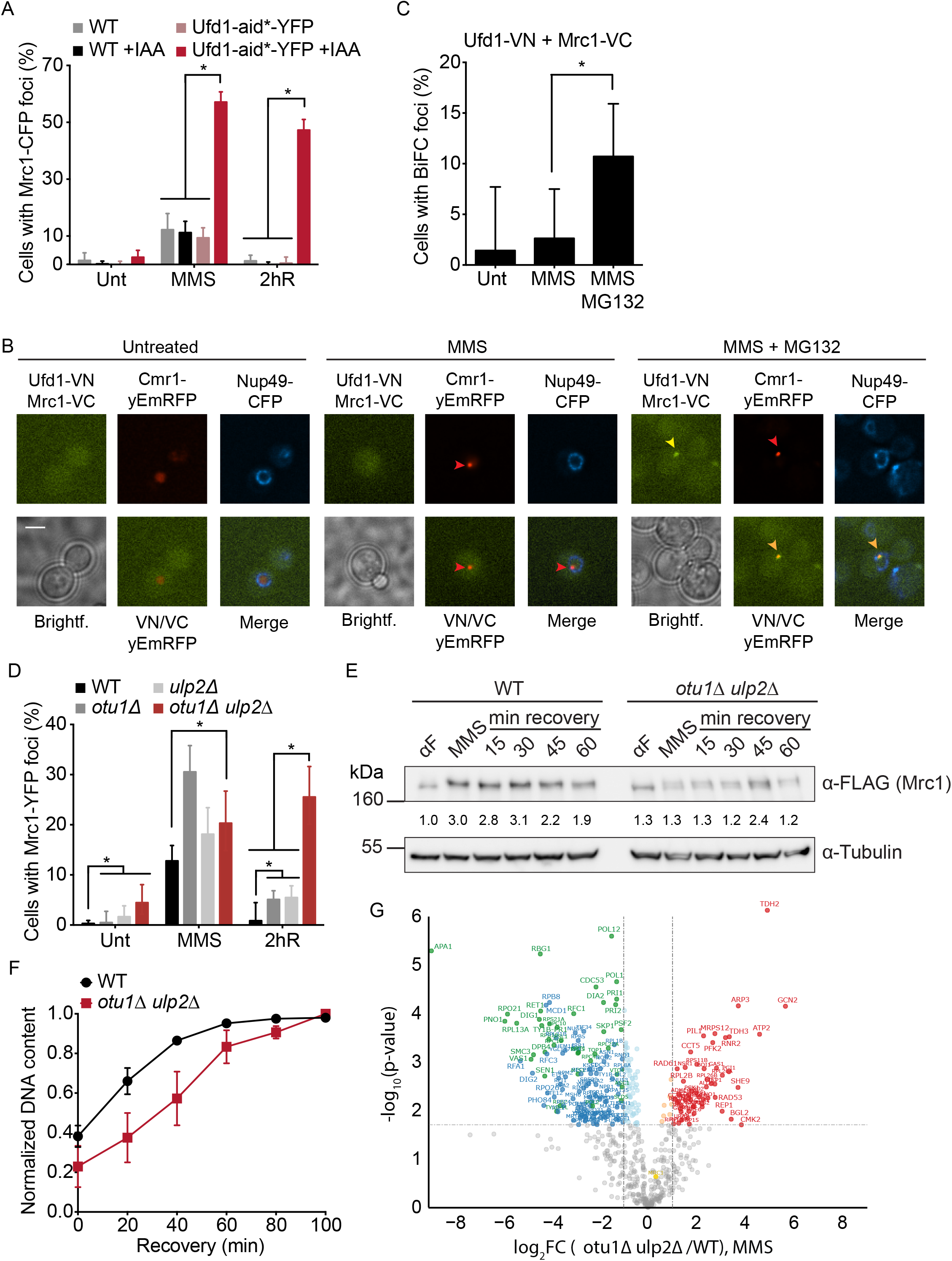
Cdc48 cofactors and Ulp2 are required for clearance of Mrc1 from INQ. **A.** Mrc1 persists in INQ in the absence of the Cdc48 cofactor Ufd1. The graph shows the percentage of asynchronous WT and Ufd1-aid*-YFP cells (CC207-9D and CC207- 3B) with or without auxin displaying Mrc1-CFP foci when untreated (Unt), treated with 0.03% MMS for 45 minutes and after 2 hours recovery from MMS (2hR). Values are averages of two experiments. Minimum 200 cells were analysed and error bars represent 95% confidence intervals. *, P < 0.0001. **B.** Mrc1 and Ufd1 interact in INQ after MMS-induced replication stress with concomitant proteasomal inhibition. Cells expressing Ufd1-VN, Mrc1-VC and Cmr1-yEmRFP (CC301-4C) were transformed with pML4 for expression of Nup49-CFP and imaged untreated (Unt) and after 2 hours 0.05% MMS treatment with or without MG132. Arrowheads point to INQ foci. Scale bar, 3 µm. **C.** Quantitation of the experiment performed in **B**. A minimum of 70 cells were analysed for each treatment. Error bars represent 95% confidence intervals. *, P < 0.05. **D.** Mrc1 persists in INQ in the absence of the de-ubiquitylation enzyme Otu1 and the SUMO de-conjugating enzyme Ulp2. The graph shows the percentage of asynchronous WT, *otu1Δ*, *ulp2Δ* and *otu1Δ ulp2Δ* cells (IG147, CC143, CC147-3B and CC181-40C) displaying Mrc1-YFP foci when untreated (Unt), treated with 0.03% MMS for 45 minutes and after 2 hours recovery from MMS (2hR). Values are averages of two to five experiments. Minimum 200 cells were counted and error bars represent 95% confidence intervals. *, P < 0.05. **E.** Mrc1 protein level is reduced after replication stress in the absence of *OTU1* and *ULP2*. Whole cell extracts were prepared from WT or *otu1Δ ulp2Δ* cells expressing Mrc1-FLAG (CC201-7B and CC214-2B) harvested after alpha factor arrest, 45 min after release into 0.03% MMS and after 15, 30, 45, and 60 minutes recovery from MMS. Mrc1-FLAG was analysed by immunoblotting. Tubulin was used as a loading control. Tubulin-normalized Mrc1-FLAG protein levels relative to the sample harvested after alpha factor arrest of WT cells are indicated below the blot. **F.** Replication recovery after MMS-induced replication stress is impaired in the absence of *OTU1* and *ULP2*. WT and *otu1Δ ulp2Δ* cells (CC44-8A and CC181-17C) were harvested before and after alpha factor arrest, after 45 minutes release into 0.03% MMS and after 20, 40, 60, 80, and 100 minutes recovery from MMS in medium containing nocodazole. The graph shows the normalized DNA content as measured from the flow cytometry data in samples taken immediately before and at indicated time points after recovery from MMS. Values are averages of 8 and two experiments, respectively, and error bars represent SEM. **G.** Proteomics analysis of Mrc1 interacting proteins in replication stressed *otu1*Δ *ulp2*Δ cells. WT and *otu1*Δ *ulp2*Δ cells (ML1168-1B and ML1165-5D) were arrested in G1 phase using alpha-factor and released into S phase with 0.03% MMS. Cells were harvested for protein extraction 45 min after release from alpha-factor arrest. After immunoprecipitation of Mrc1, co- immunoprecipitating proteins were identified by LC-MS/MS. Proteins that display a significantly reduced interaction with Mrc1 in MMS-treated *otu1*Δ *ulp2*Δ cells relative to MMS-treated wild-type cells are highlighted in green and blue, depending on whether or not the protein was already identified as a significant interactor of Mrc1 in wild-type cells under the same condition (Table S6), respectively. Proteins that display a significantly increased interaction with Mrc1 in MMS-treated *otu1*Δ *ulp2*Δ cells relative to MMS-treated wild-type cells are highlighted in green and red, depending on whether or not the protein was already identified as a significant interactor of Mrc1 in wild-type cells under the same condition (Table S6), respectively. Dashed lines indicate thresholds (-1 > log2FC (*otu1Δ ulp2*Δ/WT) > 1; -log10(p-value) for the protein with the highest significant posthoc adjusted p-value below 0.05).

To test if Cdc48 cofactors colocalize with Mrc1 foci, MMS-treated cells expressing Mrc1-YFP and Otu1-, Ubx5- or Wss1-yEmRFP or Mrc1-CFP and Ufd1-aid*-YFP were imaged. None of the four Cdc48 cofactors formed foci in response to MMS (Table S1), suggesting that they do not accumulate in INQ upon MMS-induced replication stress. However, this could be due to the localization being highly dynamic and transient. We therefore turned to BiFC, which can capture transient interactions and reveal their site of interaction in the cell (Shyu & Hu, 2008). To this end, cells expressing Otu1-, Ubx5- or Ufd1-VN/VC and Mrc1-VC/VN were imaged 2 hours after MMS and/or MG132 treatment. In a subset of cells, a focal Venus fluorescence signal was detected, indicating an interaction of Mrc1 with these cofactors in INQ as demonstrated by the colocalization with the INQ marker Cmr1 (Figures 4B, 4C, S6A and Table S2). Notably, pan-nuclear Venus fluorescence signals were observed in cells expressing pairwise combinations of Otu1-, Ubx5- and Ufd1-VN/VC (Figure S6B and Table S2), suggesting that these Cdc48 cofactors can associate with the same Cdc48 hexamer. Taken together these data indicate that Mrc1 interacts transiently with Cdc48 cofactors Otu1, Ubx5 and Ufd1 in response to replication stress and proteasomal inhibition.

Ufd1 recognizes both ubiquitylated and SUMOylated substrates (Meyer *et al*., 2002; Nie *et al*, 2012) and it is therefore possible that lack of ubiquitylation or SUMOylation of Mrc1 could abolish the interaction between Mrc1 and Ufd1, thus leading to an inability to clear Mrc1 from INQ upon recovery from MMS-induced replication stress. Mrc1 is SUMOylated at K783 by the SUMO E3 ligase Mms21 (Stefan Jentsch and Ivan Psakhye, personal communication). *mms21-11* mutants expressing Mrc1-YFP or cells expressing mrc1-K783R-YFP, which would abolish sumoylation at K783, were therefore examined for persistence of YFP foci after 2 hours of recovery from MMS treatment. Mrc1-YFP foci did persist in *mms21-11* mutant cells (Table S1), though not to the same degree as upon auxin-induced degradation of Cdc48 or Ufd1 (Figure 3A and 4A). In contrast, when analysing another SUMO E3 deletion mutant, *siz2Δ,* persistence of Mrc1-YFP foci was not observed (Table S1), indicating that Mms21-catalysed SUMOylation could be important for Mrc1 clearance from INQ. Yet, persisting foci were not observed for mrc1-K783R-YFP (Table S1), suggesting that other lysines in the Mrc1 protein may be SUMOylated by Mms21 in the absence of K783 or the effect of mutating *MMS21* may be indirect. We attempted to identify the ubiquitylation sites in Mrc1 by several mass spectrometry approaches, but were unable to reproducibly detect specific sites, which prevented mutagenesis of Mrc1 ubiquitylation sites (data not shown). Finally, possible effects of ubiquitylation- or SUMOylation-dependent ubiquitylation events were tested by analysing the *san1Δ* ubiquitin ligase mutant and SUMO-targeted ubiquitin ligase (STUbL) deletion mutants *rad18Δ*, *slx8Δ* and *uls1Δ* for Mrc1 foci upon treatment with and recovery from MMS. While none of these enzymes were required for Mrc1 re-localization to INQ upon MMS-induced replication stress, deletion mutants of *SAN1* and *ULS1* displayed persistence of Mrc1-YFP foci in INQ (Table S1). This indicates a contribution of ubiquitylation mediated by San1 and Uls1 in the clearance of Mrc1 from INQ or increased re- localization of Mrc1 to INQ in these mutants.

Taken together, these data suggest that Mrc1 is targeted for processing by Cdc48 in INQ through recognition by the Cdc48 cofactors Otu1, Ubx5 and Ufd1, and that Mrc1 clearance from INQ at least partially depends on Mms21-mediated SUMOylation and ubiquitylation by San1 and Uls1.

### De-modification recycles Mrc1 from INQ to the replication fork during replication checkpoint recovery

It has been suggested that Cdc48 might rescue some substrates from proteasomal degradation by associating with the de-ubiquitylating enzyme (DUB) Otu1 (Baek *et al*., 2013; Jentsch & Rumpf, 2007). As we observed a return of Mrc1 to the fork upon release from MMS-induced replication stress (Figure 1G and S1) as well as a focally localized perinuclear interaction between Mrc1 and Otu1 upon MMS treatment by BiFC (Figure S6A, Table S2), we hypothesized that INQ-localized Mrc1 could be de- ubiquitylated and released from INQ in order to return to the replication fork and support efficient replication completion. If this were the case, abolishing the relevant DUB would lead to a persistence of Mrc1 in INQ during recovery from MMS-induced replication stress. Similarly, de-phosphorylation of Mrc1 by Pph3 or de-SUMOylation by Ulp1 or Ulp2 could promote recycling of Mrc1 from INQ to the replication fork. To test this hypothesis, Mrc1-YFP expressing cells deleted for *OTU1* and other de-modification enzymes were imaged after 2 hours of recovery from MMS-induced replication stress. The *pph3Δ* mutant did not display persisting Mrc1-YFP foci during recovery from MMS-induced replication stress (Table S1), indicating that either Mrc1 is not a target of Pph3 or that de-phosphorylation of Mrc1 is not imperative for release of Mrc1 from INQ. Next, the importance of de-SUMOylation for Mrc1 release from INQ was examined by eliminating the activity of the two SUMO de-conjugating enzymes Ulp1 and Ulp2. The *ulp2Δ* but not the *ulp1NΔ338* (Zhao *et al*, 2004) mutant exhibited an elevated level of persisting Mrc1-YFP foci upon recovery from MMS (Figure 4D and Table S1), suggesting a contribution of Ulp2 mediated de-SUMOylation to release Mrc1 from INQ, which is consistent with the reported interaction of Ulp2 with Cdc48 (Psakhye *et al*, 2019). Compromising de-ubiquitylation by deletion of the Cdc48- associated DUB *OTU1* also induced an elevated level of persisting Mrc1-YFP foci. Importantly, when imaging the double mutant *otu1Δ ulp2Δ*, persisting Mrc1-YFP foci were observed to the same degree as in the absence of Cdc48 (compare Figure 4D and 3A). This indicates that both de-ubiquitylation and de-SUMOylation contributes to clearance of Mrc1 from INQ during recovery from MMS-induced replication stress.

Next, the possibility of Mrc1 being a substrate of Ulp2 was addressed using BiFC to investigate whether the proteins physically interact. When cells expressing Ulp2-VN and Mrc1-VC were imaged after 2 hours of MMS treatment, a focal BiFC signal was observed in a subset of cells similar to what was found for Otu1 (Figure S6C, Table S2), suggesting that the two enzymes interact with Mrc1 in a pattern resembling INQ localization. Interestingly, a focal signal was also observed in cells expressing Otu1- and Ulp2-VN/VC (Figure S6B), indicating that the two enzymes interact with each other and thus could potentially act on Mrc1 in tandem. In addition, total Mrc1-FLAG protein level was reduced in the *otu1Δ ulp2Δ* mutant during and in recovery from MMS-induced replication stress and Mrc1 remained hyperphosphorylated (Figure 4E), indicating an increased degradation of Mrc1 in this mutant. To test if de-modification of Mrc1 by Ulp2 and Otu1 facilitates recycling of Mrc1 from INQ to the replication fork during recovery from MMS treatment, we performed immunoprecipitation of Mrc1 in *otu1Δ ulp2*Δ cells before, during and after treatment with MMS and analysed the co-immunoprecipitated proteins by mass spectrometry (MS). This analysis shows that the association of Mrc1 with replisome components is further reduced in the *otu1Δ ulp2*Δ mutant in all conditions compared to WT cells upon MMS treatment and during recovery from MMS (Figures 4G, S1 and S2C). Especially the interaction between Mrc1 and polymerase epsilon was highly attenuated (Figure S2C). The proteomics analysis of the Mrc1 interactome in the absence of Otu1 and Ulp2 also showed that the interaction of Mrc1 with the SCF-Dia2 ubiquitin ligase complex was reduced (Figures S1, 4G, S2C and S5G, and Table S7). In contrast, the interaction of Mrc1 with the Rad53 checkpoint kinase was increased (Figure S4D), which is consistent with the hyper-phosphorylation of Mrc1 (Figure 4E). Finally, absence of *OTU1* and *ULP2* showed delayed replication completion during recovery from MMS-induced replication stress when compared to WT (Figure 4F). Collectively, these data indicate that Cdc48, besides delivering Mrc1 for proteasomal degradation, can also rescue Mrc1 from proteolysis and instead promote Mrc1 de-ubiquitylation and de-SUMOylation by Otu1 and Ulp2 resulting in the release of Mrc1 from INQ and completion of DNA replication.

### Replication restart is compromised in the absence of Dia2 and Cdc48

In order to investigate whether the slow replication recovery we observed in mutants compromised for Mrc1 turnover and INQ localization dynamics is due to replication fork defects, we performed DNA combing in WT and *dia2Δ* cells as well as in Ufd1- aid*-YFP and Cdc48-aid cells in the presence of auxin. Cells synchronized in G1 were released into MMS-containing medium in the presence of iodo-deoxyuridine (IdU). After 45 minutes, cells were transferred to fresh medium without MMS for 30 minutes in the presence of chloro-deoxyuridine (CldU, Figure 5A). Following DNA combing, specific antibodies were used to stain for IdU and CldU in order to discriminate between DNA replicated in the presence of MMS or during recovery from the drug. Representative DNA fibers are shown in Figure 5B. In WT cells, the median tract length was 55.0 kb and 19.2 kb for IdU and CldU tracts, respectively (Figure 5C, 5D and S7). This corresponds to an average fork rate of 1.2 kb/min during MMS treatment and 0.64 kb/min during recovery from MMS. This is in agreement with previous findings (Luke *et al*, 2006) and shows that replication progresses at approximately 60% of the normal rate through MMS-alkylated DNA (Hodgson *et al*., 2007). Interestingly, the median IdU tract length was significantly shorter in the absence of Dia2, Ufd1, and Cdc48, with *dia2Δ* and auxin-treated Cdc48-aid cells showing the shortest median tract lengths (34.1 and 31.5 kb, respectively) (Figure 5C). This indicates that replication fork speed is further reduced during MMS treatment in these mutants. Furthermore, CldU tract lengths were significantly reduced in all three mutants compared to WT (Figure 5D), indicating either a general reduction in fork progression or a delay or inability in replication fork restart compared to WT cells. Since replication kinetics were similar to WT in unchallenged *dia2Δ* and auxin-treated Cdc48-aid cells (Figure S3E), we considered the second hypothesis more plausible. Therefore, we directly assayed fork restart from the DNA combing data by calculating the percentage of IdU-incorporating forks that exhibit continued CldU-incorporation during recovery from MMS (Figure 5E). While 55% of forks had restarted after 30 minutes of recovery from MMS treatment in WT cells, only 33%, 35% and 17% of forks had restarted in *dia2Δ* and auxin-treated Ufd1-aid*-YFP or Cdc48-aid cells, respectively. This suggests that fork restart is compromised in the absence of Dia2 and Cdc48 thus leading to the delayed completion of replication as observed by flow cytometry (Figures S3D and 3D). This is further corroborated by the increased asymmetry observed in bidirectional CldU tracts (sister forks) in *dia2Δ* cells and auxin-treated Cdc48-aid cells (Figure 5F), suggesting an increased prevalence of stalled or collapsed forks in these mutants. Notably, Ufd1- depleted cells exhibited the mildest defect in replication recovery by DNA combing and no significant defect by flow cytometry analysis (Figure 5F and S5B). This suggests that there could be redundancy between Ufd1 and other Cdc48 cofactors allowing for sufficient Mrc1 to be recycled to the fork for replication completion. Taken together, these data suggest that de-modification and release of Mrc1 from INQ supports replication completion during checkpoint recovery by promoting replication fork restart.

**Figure 5.**
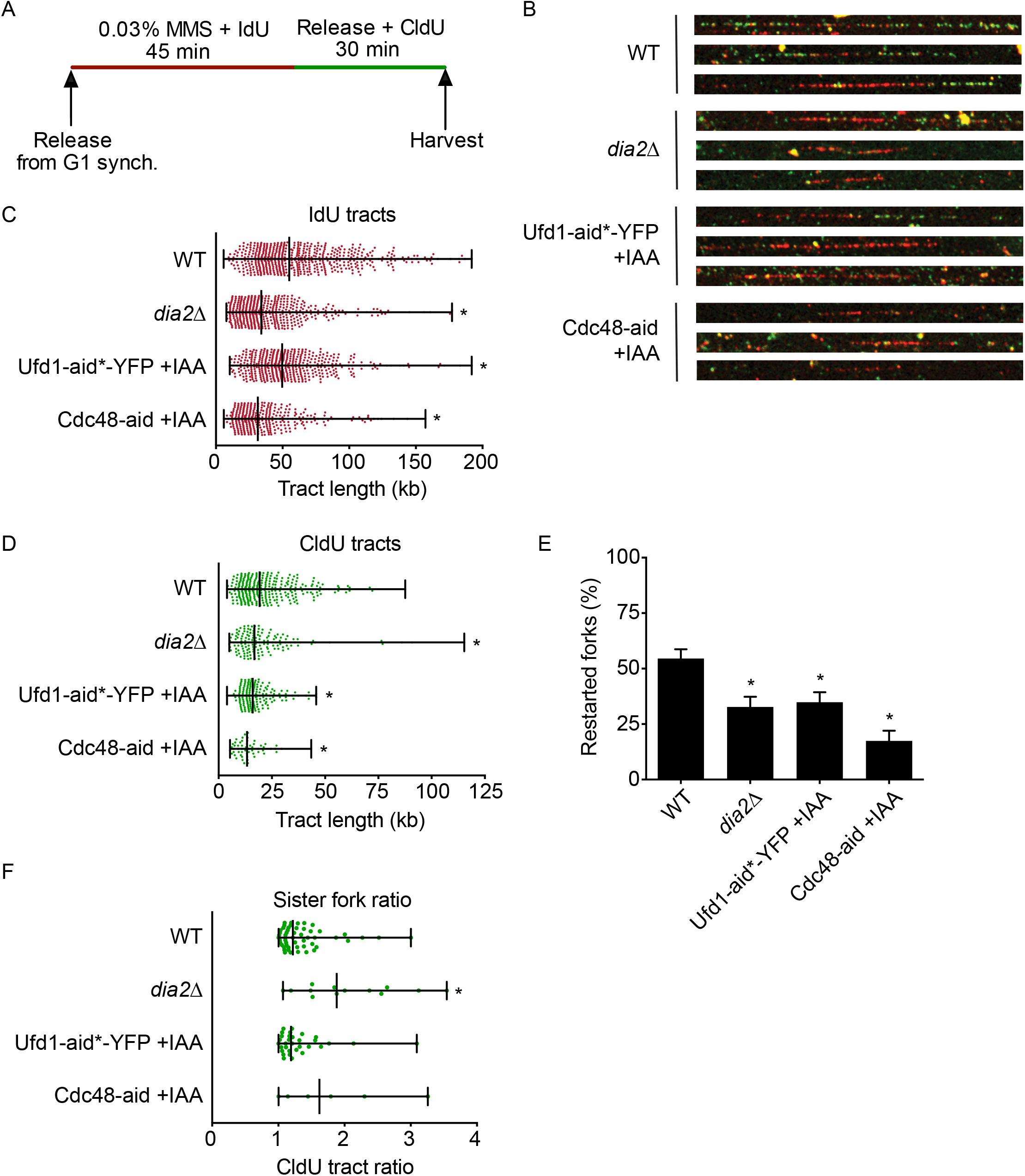
Cdc48-Ufd1 and Dia2 are required for replication restart following replication stress. A-F. DNA combing demonstrates defective restart of replication forks in *dia2Δ*, Ufd1-aid*-YFP, and Cdc48-aid mutants. **A.** Schematic representation of the experimental setup. WT, *dia2Δ,* Ufd1-aid*-YFP, and Cdc48-aid cells (ML929- 3C, CC231-1B, CC230-38D, ML941-2A) were grown to exponential phase and synchronized by addition of alpha factor. Ufd1-aid*-YFP and Cdc48-aid cells were synchronized in the presence of IAA to induce degradation of the proteins. Cells were released for 45 minutes into 0.03% MMS in the presence of IdU and then allowed to recover for 30 minutes in the presence of CldU in medium without MMS. Finally, samples were harvested, processed for DNA combing and stained for IdU (red) and CldU (green). The experiment was performed twice and the measurements pooled. **B.** Representative images of DNA fibers. **C-F.** IdU tract length (**C**), CldU tract length (**D**) and sister fork ratios (**F**) are shown as scatter plots with the minimum, median and maximum values indicated. Sister fork ratios are calculated as the length of the longer divided by the length of the shorter CldU tract. *, P < 0.05. **E.** Percentage of restarted forks. Error bars represent 95% confidence intervals. *, P < 0.0001.

## Discussion

In this study, we dissect the shuttling of Mrc1 between the replication fork and INQ during DNA replication stress and provide evidence that this response is required for the efficient restart of DNA replication during recovery from stress. We propose the following model to explain our results (Figure 6): During MMS-induced replication stress, Mrc1 dissociates from the replication fork (Figure 1C-D). Next, Dia2 drives the sequestration of Mrc1 into INQ by promoting Btn2 accumulation (Figure 2H) and likely also by ubiquitylation of Mrc1 (Mimura *et al*., 2009). Further, the localization of Mrc1 to INQ appears to be promoted by sumoylation, because Mrc1 accumulates in INQ upon deletion of the SUMO-deconjugating enzyme Ulp2 (Figure 4D). However, sumoylation of Mrc1 at K783 (Stefan Jentsch and Ivan Psakhye, personal communication) is not required for its localization to INQ, which indicates that other sites or targets could be sumoylated as well. In INQ, ubiquitylated and sumoylated Mrc1 is recognized by the Cdc48 cofactor Ufd1 leading Cdc48 to process Mrc1 to either degradation by the proteasome or Otu1- and Ulp2-mediated de-modification and release from INQ, facilitating replication checkpoint recovery. Since the Ufd1 depletion only induces a mild replication restart defect (Figure 5F and S5B), we conclude that de-modification of Mrc1 by Otu1 and Ulp2 are more important for replication stress recovery than Ufd1 or that other Cdc48 cofactors can act redundantly with Ufd1 in targeting Mrc1 to the proteasome. During recovery, Mrc1 returns to the replication fork in order to provide support for efficient restart and completion of DNA replication.

**Figure 6.**
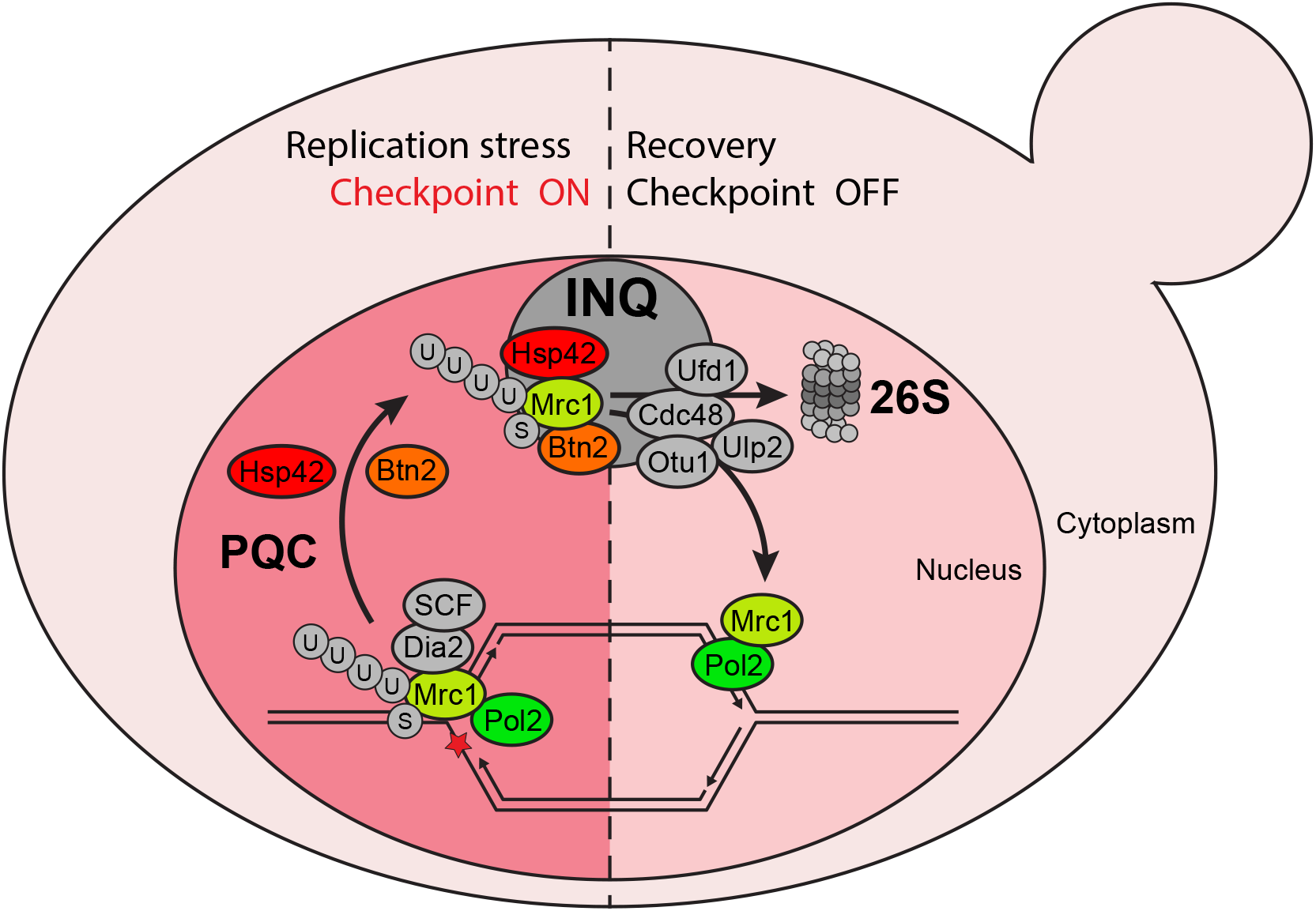
Shuttling of Mrc1 between the replication fork and INQ influences Mrc1 protein turnover and replication stress recovery. Model for the mechanism and functional impact of Mrc1 re-localization during and in recovery from MMS-induced replication stress. Mrc1 is phosphorylated, ubiquitylated and potentially SUMOylated during replication stress leading to its dissociation from the replication fork, where it is recognized by the nuclear protein quality control (PQC) system. Upon saturation of the PQC system, the Hsp42 and Btn2 chaperones sequester PQC substrates in INQ. Cdc48 extracts Mrc1 from INQ using the Ufd1 cofactor to promote degradation by the 26S proteasome, or using the cofactor Otu1 or Ulp2 to facilitate de-ubiquitylation and de-sumoylation, respectively. De-modified Mrc1 returns to the replication fork to complete DNA replication or engage in repair activities.

Our finding that ectopic expression of Btn2 in the absence of genotoxic stress is sufficient to sequester Mrc1 in INQ points to Btn2 as a central factor in the formation of INQ. The role of Btn2 in regulating Mrc1 may extend to other types of stress since Btn2 expression is also induced by ethanol (Yamauchi & Izawa, 2016) and since Mrc1 localization was shown to be regulated by several types of stress, including ethanol (Duch *et al*, 2018; Voordeckers *et al*, 2020). Possibly, Btn2 expression is promoted by Dia2 (Figure 2H) as a consequence of the ability of Dia2 to control assembly of the RSC chromatin remodeling complex and facilitate inducible gene expression (Andress *et al*., 2011). Moreover, upon activation of the replication checkpoint, Dia2 is stabilized (Kile & Koepp, 2010; Mimura *et al*., 2009) and consequently, Btn2 may accumulate further. Sequestration of Mrc1 in INQ in the absence of genotoxic stress is also observed when cells are treated with the proteasome inhibitor MG132 (Gallina *et al*., 2015). This may be related to Mrc1 being mostly an intrinsically disordered protein (Meszaros *et al*., 2018). Hence, Mrc1 may be recognized as a misfolded protein and sequestered by the Btn2 chaperone (Gallina *et al*., 2015; Miller *et al*., 2015) upon Btn2 expression. Importantly, deletion of *BTN2* in cells defective of Cdc48-mediated Mrc1 turnover, further delays replication recovery (Figure 3D), indicating that recovery from MMS-induced replication stress is supported by sequestering Mrc1 in INQ as well as Cdc48-mediated degradation of Mrc1.

Mrc1 is not the only INQ-localized protein regulated by Cdc48. Cmr1 and Pph3 both localize to INQ upon MMS-induced replication stress and proteasomal inhibition (Gallina *et al*., 2015) and both persist in INQ in the absence of Cdc48 (Figure S4A). This indicates that Cdc48 has a more general function in dissolving INQ. As we have previously shown the presence of SUMO in INQ (Gallina *et al*., 2015), this dissolving activity could be analogous to that suggested for the Cdc48-mediated dissolving of DNA repair foci held together by interactions between SUMO and SUMO-interacting motifs (SIMs) (Jentsch & Psakhye, 2013; Psakhye & Jentsch, 2012). Interestingly, a recent study into nuclear protein quality control, identified Cdc48 as a regulator of misfolded, San1-ubiquitylated substrates (Gallagher *et al*, 2014). San1 is the primary ubiquitin ligase responsible for targeting nuclear misfolded proteins for proteasomal degradation and in the absence of functional Cdc48, San1 substrates become more insoluble and persist in inclusion bodies in the nucleus (Gallagher *et al*., 2014; Gardner *et al*, 2005). Several parallels exist between these observations and INQ. Firstly, misfolded proteins localize to INQ (Gallina *et al*., 2015), Mrc1 persists in INQ during recovery from MMS-induced replication stress in the absence of *SAN1* (Table S1) and Mrc1-, Cmr1- and Pph3-INQ foci intensify and persist in the absence of Cdc48 (Figure 3A and S4A). These parallels lead us to suspect that the inclusions observed by Gallagher *et al*. are INQ-related (Gallagher *et al*., 2014). In order to further understand the impact of sub-nuclear localization on nuclear processes, it will be important for future studies to further uncover the function of Cdc48 in disaggregation, segregation and dissolving of sub-nuclear structures like INQ and DNA repair foci.

In addition to showing that Cdc48 is central to Mrc1 degradation after MMS-induced replication stress, we also present data suggesting that Mrc1 is recognized by three different Cdc48 cofactors: Ufd1, Ubx5 and Otu1 (Figure S6A and Table S2. Jentsch and Rumpf suggested that by associating with different substrate-recruiting and -processing cofactors, Cdc48 could act as a molecular gearbox, targeting substrates to different fates (Jentsch & Rumpf, 2007). Mrc1 may be such a substrate. Our data indicate that while Cdc48 is stably present in INQ during replication stress (Figure 3B), the Cdc48 cofactors are not (Table S1). Instead, they may dynamically associate with Cdc48 in INQ and thus direct Cdc48-dependent processing of INQ- localized substrates to different fates. In the case of Mrc1, recognition by Ufd1 most likely results in proteasomal degradation as the interaction between Mrc1 and Ufd1 was stabilized in the presence of MG132 (Figure 4B and 4C), similar to what has been shown previously for other Cdc48-Ufd1-Npl4 substrates (Bays *et al*, 2001; Franz *et al*, 2011; Raman *et al*, 2011; Verma *et al*., 2011; Ye *et al*, 2003). Less is known about Ubx5, but this cofactor has been shown to facilitate the UV-induced degradation of the RNA pol II subunit Rpb1 (Verma *et al*., 2011). The same may be true for Mrc1, as absence of either Ufd1 or Ubx5 caused Mrc1 to persist in INQ during recovery from MMS-induced replication stress (Figure 4A and Table S1). Finally, recognition by Cdc48-associated Otu1 may result in the removal of ubiquitin modifications from Mrc1 (Rumpf & Jentsch, 2006; Stein *et al*, 2014). Since Mrc1 is required for efficient replication in unperturbed cells (Gispan *et al*., 2014; Hodgson *et al*., 2007; Szyjka *et al*., 2005), de-modification of Mrc1 may rescue the protein from proteasomal degradation and facilitate Cdc48-mediated extraction of Mrc1 from INQ. De-modified Mrc1 could thus return to the replication fork during recovery from MMS-induced replication stress in order to efficiently restart and complete replication. This is supported by our observations that Mrc1 persists in INQ in the absence of Otu1 (Figure 4D), that the interaction between Mrc1 and Ctf4 is re-established upon recovery from MMS-induced replication stress (Figure 1G), and that replication completion is delayed in the absence of Otu1 and Ulp2 (Figure 4F).

As Mrc1 mediates the replication checkpoint by facilitating the recruitment of Mec1 to chromatin and the activation of Rad53 (Alcasabas *et al*., 2001; Osborn & Elledge, 2003), why would it be beneficial to sequester Mrc1 in INQ upon MMS-induced replication stress? Our finding that Cdc48 depletion resulted in a concomitant accumulation of Mrc1 in INQ and increased interaction with the Rad53 checkpoint kinase indicates that localization of Mrc1 to INQ may in fact enhance checkpoint signaling. Since Mrc1 promotes high speed replication (Gispan *et al*., 2014; Hodgson *et al*., 2007; Szyjka *et al*., 2005), dissociation of Mrc1 from the fork may at the same time reduce the speed of replication of damaged DNA or facilitate its replication by translesion polymerases (Voordeckers *et al*., 2020), which is consistent with the 4-fold increased mutation rate reported for *mrc1*Δ cells (Gallina *et al*., 2015). Alternatively, toxic fork regression promoted by Mrc1 (Menin *et al*, 2018) may be prevented by dissociation of Mrc1 from the fork. Consistently, the *mrc1*^AQ^ mutant, which associates tighter with the replication fork during stress (Lou *et al*., 2008) and exhibits reduced localization to INQ (Gallina *et al*., 2015), causes increased triplet instability (Gellon *et al*, 2019).

The proteomics analysis conducted in this study confirmed many of the known interactors of Mrc1 that can be explained by its direct interaction with the replisome (Baretic *et al*, 2020), but also identified new physical interactors of Mrc1 (Table S6).

Some of these new interactions may be bridged by other proteins, but they may nevertheless be relevant to Mrc1 function. For example, Mrc1 exhibits robust co-immunoprecipitation with the Rrm3 helicase (S5C), which mirrors the behaviour of the replisome interactions, as well as a constitutive interaction with all four subunits of casein kinase II (Cka1, Cka2, Ckb1, Ckb2) (Figure S2B and Table S6). Notably, cells lacking Mrc1 become highly dependent of Rrm3 to cope with replication stress (Schmidt & Kolodner, 2006; Szyjka *et al*., 2005) and our data now suggest a physical link between the two proteins. Relatively few interactions of Mrc1 increase with MMS treatment with some notable exceptions including the Rad5 helicase and ubiquitin ligase, all five cohesin subunits (Smc1, Smc3, Irr1, Mcd1, Pds5), and the Mms1-Rtt101 complex (Figures S4D, S5C and S5D; Table S7). In this context, it is interesting to note that Mrc1 and the Mms1-Rtt101 complex were reported to be required for establishing sister chromatid cohesion during post-replication repair (Xu *et al*, 2004; Zhang *et al*, 2017), and Mms1-Rtt101 promotes replication of damaged DNA (Zaidi *et al*, 2008). The functional implications of the Rad5-Mrc1 interaction will be the subject of future studies.

We observed major changes in the Mrc1 interactome in mutants that affect its localization to INQ. For example, *btn2*Δ further reduces the interaction with polymerase epsilon, cohesin, the Senataxin helicase (Sen1) and the regulators of transcription-coupled repair Rpb4-Rpb7 upon MMS treatment (Figures S2C and S4D; Table S7), suggesting that Btn2 could be required for Mrc1 to maintain or engage in these interactions during replication stress. To explain these changes, we speculate that Mrc1, which has dissociated from stalled replication forks due to its post-translational modification, is required to first be sequestered in INQ by Btn2 and de-modified, before it can engage in these interactions.

Depletion of Cdc48 caused a persistence of Mrc1 in INQ during recovery from MMS treatment (Figure 3A), which was accompanied by a dramatic reduction in the interaction of Mrc1 with Rad5, a helicase and ubiquitin ligase, and the Mms1-Rtt101 complex (Figures S4D and S5D), and an equally dramatic increase in the interaction with the Rad53 checkpoint kinase (Figure 3E and S4D), suggesting that checkpoint signaling is increased when Cdc48 fails to release Mrc1 from INQ. Since Mrc1 is responsible for mediating the replication stress checkpoint from the apical Mec1/ATR kinase to the Rad53 effector kinase (Osborn & Elledge, 2003) through a direct interaction with both Mec1 and Rad53 (Chen & Zhou, 2009), we predict hyper- activation of the checkpoint upon Cdc48 depletion. The interactions of Mrc1 with the chaperones Hsp26, Hsp42, Ssa2, Ssa4, and the Chaperonin Containing TCP-1, and co-chaperones Ydj1 and Sis1 were also reduced upon Cdc48 depletion (Figures 3E S1, S5F and Table S7), suggesting that Mrc1 becomes inaccessible to these chaperones despite its localization to INQ, when the Cdc48 segregase is depleted. Taken together, these changes indicate that Cdc48-mediated release of Mrc1 from INQ may be required for recognition of Mrc1 by other chaperones and for its interaction with Rad5, which promotes MMS tolerance via regression of stalled replication forks (Blastyak *et al*, 2007).

Similarly to depletion of Cdc48, disruption of the Otu1 ubiquitin and Ulp2 SUMO peptidases increased the accumulation of Mrc1 in INQ (Figure 4D). However, simultaneous loss of Otu1 and Ulp2 caused a more dramatic decrease in the interaction of Mrc1 with replisome components (especially PCNA (Pol30) and RPA), Figure S4D), factors involved in post-replication repair (Mms1-Rtt101, Figure S5D) and resolution of replication-transcription conflicts (Sen1, Figure S4D) (Appanah *et al*, 2020) than depletion of Cdc48, while chaperone interactions were largely unaffected (Figure S5F). Since completion of DNA replication after MMS treatment is similarly delayed in Cdc48 depleted and *otu1Δ ulp2*Δ mutant cells, this result suggests that the Hsp26 and Hsp42 chaperone interactions of Mrc1 could be less critical for completion of DNA replication after MMS treatment probably due to a high degree of redundancy in their substrates (Haslbeck *et al*, 2004).

The proteomics analysis of the Mrc1 interactome in mutants that either abolish or enhance localization of Mrc1 to INQ indicates that localization to INQ *per se* does not enforce specific interactions, but rather that specific chaperones and enzymatic activities (de-ubiquitylation and SUMO-deconjugation) may affect Mrc1 localization, interactions and turnover during replication stress. Given that only a fraction of Mrc1 protein in 10-15% of wild-type cells re-localize to INQ during stress, we speculate that the formation of INQ reflects a situation where the protein quality control machinery of the cell is overwhelmed by the quantity of substrate, leading to formation of INQ by liquid-liquid phase separation and/or sequestration by Btn2 and Hsp42. Under such circumstances Cdc48 aided by Ufd1, Otu1 and Ulp2 becomes important for degrading or recycling Mrc1 back to the replication fork during recovery from replication or proteotoxic stress.

Mrc1 function seems to be generally conserved in humans, as the human ortholog Claspin, like Mrc1, is required for unperturbed replication as well as for mediating the ATR-dependent activation of the replication checkpoint effector kinase Chk1 (Chini & Chen, 2003, 2004; Petermann *et al*, 2008). In addition, replication checkpoint-dependent phosphorylation is also required for Claspin function (Chini & Chen, 2006). In line with our data on Mrc1, Claspin is also ubiquitylated and targeted for degradation in response to replication stress and DNA damage by the SCF-βTrCP ubiquitin ligase, facilitating checkpoint recovery (Mailand *et al*, 2006; Peschiaroli *et al*, 2006). Finally, ubiquitylation and degradation of Claspin is also counteracted by DUBs during the response to replication stress (Martin *et al*, 2015; McGarry *et al*, 2016). There is currently no evidence, however, of an involvement of the human Cdc48 ortholog, p97, in the regulation of Claspin degradation or localization. Whether Cdc48 function with regards to Mrc1 and INQ is conserved in human cells will be a question for future studies.

## Acknowledgements

We thank Jeff Bachant, Rey-Huei Chen, John Diffley, Stephen Elledge, Stefan Jentsch, Ivan Psakhye, Hannah Klein, Michael Knop, Karim Labib, Rodney Rothstein, Helle Ulrich and Xiaolan Zhao for sharing information, strains, plasmids and reagents. ML was supported by Danish Council for Independent Research (DFF - 7014-00176) and the Villum Foundation (research grant 11407). Mass spectrometry and data analyses were performed by the Proteomics Research Infrastructure (PRI) at the University of Copenhagen (UCPH), supported by the Novo Nordisk Foundation (NNF) (grant agreement number NNF19SA0059305).

## Author contributions

M.L. and C.C. conceived the project. C.C. designed experiments with supervision from M.L. and B. P., and C.C. and J.A. performed experiments with assistance from B.P. and M.L., C.C. and J.A. performed analysis with assistance from M.L., and C.C. and M.L. wrote the manuscript. All authors edited the manuscript.

## Materials and methods

### Yeast strains and cell culture

Unless otherwise stated, yeast strains used in this study are *RAD5* derivatives of W303 (Table S3). Standard genetic techniques were used to manipulate yeast strains and standard media were used throughout this study (Sherman, 1986).

Cell cultures were grown to exponential phase before the start of experiments. G1 synchronization was achieved by addition of alpha factor (TAG Copenhagen) to a final concentration of 7.5 µg/mL and incubating cell cultures for one hour before addition of a further 3.75 µg/mL alpha factor. Incubation was continued until at least 90% of cells were in G1 phase as determined by microscopic inspection. For auxin-induced degradation of AID-tagged proteins, Indole-3-acetic acid (IAA, Sigma) was added to a final concentration of 500 µM 1.5 hours prior to start of experiments (either during alpha factor synchronization or asynchronous growth when cells had reached exponential phase). Methyl methanesulfonate (MMS, Sigma) was added to final concentrations of 0.03% or 0.05% as indicated. Were indicated, MG132 (Selleckchem, cat.no. S2619) and Nocodazole (Sigma) were added to final concentrations of 75 µg/mL and 7.5 µg/mL, respectively.

### Yeast constructs and plasmids

YFP-tagged Pph3 and Cdc48 were constructed using adaptamer-mediated PCR as previously described (Reid *et al*, 2002). Essentially, the same technique was used to construct the strains containing Mrc1-YFP and yEmRFP-tagged Otu1, Ubx5 or Wss1. In brief, the homology arms for targeted integration were generated from template genomic DNA using primers Otu1/Ubx5/Wss1-F, -end-R, -RFPdown-F and –downR. The two homology arms were fused by PCR to sequences containing *yEmRFP* joined to either the 5’- or 3’-end of *K.l. URA3* that were PCR amplified from pNEB30 or pNEB31 (Silva *et al*, 2012) using primers Cherry.Fw and 3’-int or 5’-int and yEmRFP-R. The two fusion products were co-transformed into IG147 and selected on SC-Ura. After pop-out of the *K.l. URA3* marker and selection on 5-FOA, the genomic *OTU1*, *UBX5* or *WSS1* loci were sequenced to confirm correct integration.

For generation of the *mrc1K783R* allele, primers (mrc1-K783R-R and –F) were designed to contain the mutation and a diagnostic BamHI site. These primers were used with either mrc1-K783R-A or –B to amplify the region of *MRC1* in question from template genomic DNA. Next, the resulting products were fused by PCR to sequences containing an adaptamer A sequence joined to *3’-K.l. URA3* or an adaptamer B sequence joined to *5’-K.l. URA3* which were amplified from pWJ716 (Erdeniz *et al*, 1997) using primers 5’-int and adaptamer a or adaptamer b and 3’-int. The resulting two fusion PCR products were co-transformed into ML8-9A and IG147 and selected on SC-Ura. After pop-out of the *K.l. URA3* marker and selection on 5-FOA, the genomic *MRC1* locus was amplified by PCR, digested with BamHI and sequenced to confirm presence of the mutation.

FLAG-tagged Mrc1 was created by amplification of the 3xFLAG::HIS3MX cassette of pBP81 using Mrc1-S3-F and Mrc1-S2-R primers adapted with overhangs to target the integration of the construct at the C-terminal end of *MRC1*. The PCR product was transformed into yeast and selected on SC-His. Integration was confirmed by PCR amplification and sequencing of the locus. Similarly, *BTN2-6xHA* was generated by amplification of the 6xHA::natNT2 cassette of pYM17 (Janke *et al*, 2004) using primers Btn2-S3-F and Btn2-S2-R containing *BTN2*-specific overhangs and transformation into yeast. Correct integration was confirmed by sequencing.

VN- and VC-constructs were generated by PCR amplification of the VN173::K.l. URA3 cassette of pFA6a-VN-KlURA3 (Sung *et al*, 2013) or the VC155::NatMX6 cassette of pML103 (Silva, 2016) using primers F2 and R1 followed by transformation into yeast. Correct integration was confirmed by sequencing. VN- and VC-constructs were crossed in all possible combinations and sporulated to obtain the haploid strains used in this study (for VN- and VC-tagging of the same protein, diploid strains were used).

To obtain a plasmid for AID*-YFP tagging, YFP was amplified by PCR from pWJ1164 (Lisby *et al*, 2001) using primers SmaI-XFP-F and BglII-XFP-R. Next, the resulting PCR product and the pHyg-AID*-GFP (Morawska & Ulrich, 2013) target plasmid were digested with SmaI and BglII and the plasmid fragment containing the AID* sequence and Hyg cassette was ligated to the digested YFP sequence generating pOQR3. This plasmid was used as template to amplify the AID*- YFP::hphNT1 cassette using primers Ufd1-S3-F and Ufd1-S2-R adapted with overhangs targeting integration of the product to the C-terminus of *UFD1*. Finally, the resulting PCR product was transformed into a strain containing ADH1-OsTIR1-9myc to generate the auxin degradation inducible Ufd1-aid*-YFP strain. Correct integration was confirmed by sequencing.

Gene deletions were generated by PCR amplification of the locus-specific KanMX cassette from template genomic DNA obtained from the relevant strains of the gene disruption collection (Invitrogen) using primers annealing upstream and downstream of the genomic loci and subsequent transformation into the W303 background. Gene knockouts were verified by PCR. Similarly, Mrc1-4myc::his5+ was amplified from template genomic DNA from strain Y1134 (Alcasabas *et al*., 2001) using primers annealing upstream and downstream of the *MRC1* locus, followed by transformation of the PCR product into the *RAD5* W303 background. The knock-in was verified by sequencing.

Plasmids are described in Table S4 and primer sequences are listed in Table S5.

### Live cell fluorescence microscopy

For live cell fluorescence microscopy experiments, cells were grown in synthetic complete medium supplemented with 100 µg/mL adenine at 25°C. Diploid cells were grown in SC-Lys/Trp+Ade. Fluorophores were visualized on a DeltaVision Elite microscope (Applied Precision, Inc.). Images were deconvolved and fluorescence intensities measured using Volocity software (PerkinElmer). Representation of fluorescence microscopy data in graphs was done in Prism (GraphPad software, Inc.). Statistically significant differences observed between cell populations were determined by Fisher’s exact test and differences between time courses of ratios of fluorescence intensities were determined by two-way ANOVA.

### Mrc1-FLAG immunoprecipitation

Cells were grown in YPD at 30°C and 300-400 ODs were harvested by cold centrifugation, washed in cold 1M sorbitol, 25 mM HEPES pH 7.6 and resuspended in 1 mL cold lysis buffer (100 mM HEPES pH 7.6, 200 mM KOAc, 0.1% NP-40, 10% glycerol, 10 mM NaF, 20 mM β-glycerophosphate) supplemented with 2 mM β-mercaptoethanol, 2.67 µg/mL aprotinin, 1 mM benzamidine, 0.1 mM phenylmethylsulfonyl fluoride (PMSF), 10 µg/mL leupeptin, 0.1 µM okadaic acid and 1 µg/mL pepstatin. Flash-frozen drops were prepared from the cell suspension in liquid nitrogen and processed into powder by grinding in liquid nitrogen in a freezer mill. Cell extracts were cleared by centrifugation and 20 µl transferred to 20 µl 2x Lämmli buffer and boiled for 10 minutes (inputs). 40 µl dry volume Flag-M2-agarose beads (Sigma) were equilibrated in lysis buffer + β-mercaptoethanol and added to the remaining extract prior to 1 hour incubation at 4°C to allow binding of Mrc1-FLAG and associated proteins to the beads. Extracts were washed six times in lysis buffer + β-mercaptoethanol, resuspended in 40 µl lysis buffer + β-mercaptoethanol and 0.5 mg/mL 3xFLAG-peptide (Sigma) and incubated 30 minutes at 4°C to dissociate proteins bound to the FLAG-beads. Immunoprecipitated proteins were eluted from the FLAG-beads by addition of the FLAG-bead suspension to homemade glass bead columns followed by centrifugation. FLAG-beads were washed once in 40 µl lysis buffer + β-mercaptoethanol and 0.5 mg/mL 3xFLAG-peptide, incubated 5 minutes on ice, centrifuged and the eluates combined. Immunoprecipitated proteins were isolated from eluates by TCA extraction and analysed together with input samples by immunoblotting (see below). Cell cycle stage of samples as well as arrest and release efficiency were determined by propidium iodide (PI) staining and flow cytometry (see below).

### Protein analysis

For analysis of Mrc1 protein levels, cells were grown in YPD at 30°C. 1 OD was harvested by centrifugation and snap freezing in liquid nitrogen. Whole cell extracts (WCE) were obtained by TCA extraction. Briefly, cell pellets were resuspended in 1 mL water, added 150 µl 1.85 M NaOH, 7.5% β-mercaptoethanol and incubated 15 minutes on ice to lyse cells. Proteins were precipitated by addition of 150 µl 55% TCA, further 10 minutes incubation on ice and centrifugation. Precipitated proteins were resuspended in 50 µl HU buffer (8 M urea, 5% SDS, 200 mM Tris pH 6.8, 1.5% dithiothreitol (DTT), bromphenolblue) and neutralized by addition of 2 M Tris base. Samples were heated 10 minutes at 65°C, centrifuged and the supernatant loaded on polyacrylamide gels.

Mrc1-FLAG in immunoprecipitation and WCE samples was detected by rabbit anti-FLAG (1:3000, F7425, Sigma) followed by horseradish peroxidase (HRP)- conjugated swine anti-rabbit (1:5000) while Mrc1-FLAG in input samples was detected by HRP-conjugated rabbit anti-FLAG (1:3000, A8592, Sigma). Rabbit anti-Ctf4 (1:2000, in house) followed by HRP-conjugated swine anti-rabbit (1:5000) was used to detect Ctf4. Tubulin was detected using rat anti-Tubulin (1:5000, ab6160, abcam) followed by HRP-conjugated rabbit anti-rat (1:5000, P0450, Dako). For detection of Btn2-HA, mouse anti-HA (1:2000, MMS-101R, Covance) followed by HRP-conjugated swine anti-mouse (1:5000) was used. Goat anti-Mcm2 (1:750, sc-6680, Santa Cruz Biotechnology) followed by HRP-conjugated swine anti-goat (1:5000) was used to detect Mcm2. For detection of Cdc45, rabbit anti-Cdc45 (1:2000, serum (On *et al*, 2014)) followed by HRP-conjugated swine anti-rabbit (1:5000) was used.

For quantitation of Mrc1-FLAG protein levels, blots were developed using a CCD camera (Imagequant, LAS 4000, GE healthcare) and quantitation performed in Fiji (Schindelin *et al*, 2012).

### Harvest and preparation of samples for mass spectrometry

For MS analysis (Reusswig *et al*., 2021), cells were grown in YPD at 30°C to OD600 = 0.5, G1-arrest was performed by addition of alpha-factor to a concentration of 5 µg/mL and the cells were incubated for 1 hour after which additional alpha-factor was added to a final concentration of 7.5 µg/mL. The cultures were further incubated until >90% of cells had reached G1 arrest as determined by microscopy. The arrested cells were released into fresh YPD medium with or without 0.03% MMS. 300 ODs of cells were metabolically stopped by addition of 0.1% NaN3 (final concentration) 25 min (without MMS) or 45 min (with MMS) after release from alpha-factor arrest. Similarly, 300 ODs of cells that had been incubated in MMS for 45 min and allowed to recover for 30 min without MMS before addition of NaN3 were collected. In parallel, small aliquots of cells were harvested at each time point and used to confirm cell cycle arrest and to monitor cell cycle progression on a MACSQuant or a BD FACSJazz flow cytometer (see below). Before protein extraction, cells were first washed with precooled 10 mM HEPES-KOH pH 7.9 and then with precooled lysis buffer (100 mM HEPES-KOH, 50 mM KAc, 10 mM MgAc2, 2 mM EDTA, 20 mM NaF, 10 mM *β*-glycerophosphate, 1 mM DTT). The supernatant was aspirated, and the pellets weighed and resuspended in 3 volumes (1 g ∼ 1 mL) of freshly prepared precooled lysis buffer with protease inhibitors (cOmplete protease inhibitor, PMSF, benzamidine, aprotinin, leupeptin, pepstatin, okadaic acid). The resuspended cells were snap-frozen by dripping the cell suspension into liquid nitrogen and stored at -80°C. Cells were lysed using either a Spex SamplePrep 6870 freezer/mill or by hand milling samples in liquid nitrogen with a mortar and pestle. All samples were stored at -80°C until further use.

The milled cell powder was thawed, weighed, and 0.25 volumes of freshly prepared pre-cooled glycerol buffer (100 mM HEPES-KOH pH 7.9, 300 mM KAc, 10 mM MgAc2, 2 mM EDTA, 50% glycerol, 0.5% NP-40) with inhibitors (cOmplete protease inhibitor and 1 mM PMSF) were added. The extracts were digested with 800 U of benzonase (Merck, E1014) per mL of extract for 30 min at 4°C. The supernatants of the extracts were separated from cell debris by centrifugation at 20.000g for 20 min in a precooled Eppendorf centrifuge. Next, the supernatants were incubated with anti-GFP antibody (Roche, cat.no. 11814460001) coupled to protein G Dynabeads (ThermoFisher Scientific, cat.no. 10004D) for one hour rotating at 4°C. After the incubation, the beads were washed three times with IP wash buffer (100 mM HEPES-KOH, 100 mM KAc, 10 mM MgAC2, 2 mM EDTA, 10% glycerol) containing 0.1% NP- 40 and then twice in IP wash buffer without NP-40. The beads were then transferred to new Eppendorf tubes and washed a final time in IP wash buffer without NP-40. Samples were temporarily stored at -20°C prior to MS analysis. Three biological replicates were harvested for the MS analysis.

### Mass spectrometry analysis

The IP beads were incubated at 37°C for 30 min with elution buffer 1 (2 M urea, 50 mM Tris-HCl pH 8, 2 mM DTT, 10 µg/ml trypsin) followed by a second elution for 5 min with elution buffer 2 (2 M urea, 50 mM Tris-HCl pH 7.5, 10 mM chloroacetamide). The eluates were combined and further incubated at room temperature over-night. Tryptic peptide mixtures were acidified to 1% TFA and loaded on Evotips (Evosep).

Peptides were separated on 15 cm, 150 μM ID columns packed with C18 beads (1.9 μm) (Pepsep) on an Evosep ONE HPLC applying the ‘30 samples per day’ method, and injected via a CaptiveSpray source and 10 μm emitter into a timsTOF pro mass spectrometer (Bruker) run in PASEF mode (Meier *et al*, 2018).

### LC-MS/MS data analysis

Raw MS data were analyzed with MaxQuant (v1.6.15.0). Peak lists were searched against the yeast Uniprot FASTA database combined with 262 common contaminants by the integrated Andromeda search engine. False discovery rate was set to 1% for both peptides (minimum length of 7 amino acids) and proteins. “Match between runs” (MBR) was enabled with a Match time window of 0.7 min, and a Match ion mobility window of 0.05 min. Relative protein amounts were determined by the MaxLFQ algorithm with a minimum ratio count of two.

All statistical analysis of Label-Free Quantification (LFQ) derived protein expression data was performed using the automated analysis pipeline of the Clinical Knowledge Graph (Santos *et al*, 2020). Protein entries referring to potential contaminants, proteins identified by matches to the decoy reverse database, and proteins identified only by modified sites, were removed. LFQ intensity values were normalized by log2 transformation and proteins with less than two valid values in at least one group were filtered out. The remaining missing values were imputed using a two-step mixed imputation approach. In the first step, per each group missing values of proteins with at least 60% of valid values in that group were imputed with k-nearest neighbors (kNN). In the second step, all remaining missing values were imputed with the MinProb method (random draws from a Gaussian distribution; width = 0.3 and downshift = 1.8) (Lazar *et al*, 2016).

Differentially immunoprecipitated proteins in each group comparison were identified by one-way Anova test with Benjamini-Hochberg correction for multiple hypothesis, followed by posthoc pairwise comparison t-tests using the same parameters and Benjamini-Hochberg correction.

Significantly regulated proteins are coloured in red and blue in the volcano plots for up and downregulated hits, respectively, while the proteins of interest (present in the predefined interactome) are coloured in green.

To define the interactome of Mrc1 (Table S6), we ran differential immunoprecipitation analysis for groups "wt" and "no tag" (alpha=0.01, FC(fold change)=2). The interactome for each group "untreated", "mms" and "recovery" was defined as the collection of proteins significantly more immunoprecipitated in each comparison : untreated-wt versus untreated-no tag, mms-wt versus mms-no tag or recovery-wt versus recovery-no tag. This eliminated non-specific interactions such as Rps2 (Figure S2A). Proteins were considered significantly regulated if cutoff < alpha and log2FC >= cutoff in the statistical test. For each of the other cell lines (’cdc48 depletion’, *’btn2*Δ’, *’otu1Δ ulp2*Δ’, *’apj1Δ hsp104*Δ’) we performed differential immunoprecipitation analysis (alpha=0.05, FC=2) and considered of interest, statistically significantly immunoprecipitated proteins in each pairwise comparison that was also present in the predefined interactome. In the volcano plots, statistically significantly immunoprecipitated proteins are coloured in blue (down-regulated) and in red (up-regulated), while the proteins of interest (present in the predefined interactome) are coloured in green.

### Flow cytometry

Cells were grown in YPD at 30°C and 1 mL cell culture per sample was harvested by centrifugation and fixed in 50 mM Tris pH 7.8, 70% ethanol for a minimum of 30 minutes or overnight at 4°C. Fixed cells were washed once in 50 mM Tris pH 7.8 before incubation at 37°C in 500 µl 50 mM Tris pH 7.8 + 20 µl 10 mg/mL RNase A (Sigma), 10 mM Tris pH 7.5, 10 mM MgCl2 for a minimum of 4 hours or overnight. Cells were then resuspended in 200 µl 50 mM Tris pH 7.8 + 20 µl 20 mg/mL proteinase K (Sigma), 50 % glycerol, 10 mM Tris pH 7.5, 25 mM CaCl2 and incubated 30 minutes at 50°C. Following resuspension in 500 µl 50 mM Tris pH 7.8, cells were either stored at 4°C or directly stained with PI (Sigma). To this end, cells were sonicated prior to incubation for 30 minutes at room temperature in the dark in 50 mM Tris pH 7.8 + 3 µg/mL PI. Sample PI profiles were recorded using a FACSJazz cell sorter (BD Biosciences) and 30.000 cells were measured for each sample. Data were processed using FlowJo software (v10.0.6, Tree Star, Inc.). Representation of flow cytometry data in graphs was done in Prism (GraphPad software, Inc.) and normalized DNA content was calculated as:

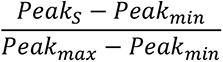

Where *Peak* refers to the most prevalent PI intensity in the flow cytometry histogram for the time point corresponding to the sample (S), asynchronous G1 or alpha factor sample (min) and asynchronous G2 or 100 minute release sample (max) for each strain.

### DNA combing

Cells were grown in YPD at 30°C until exponential phase and synchronized in G1 by addition of alpha factor. Cells were washed and released into S phase in medium containing 0.03% MMS and 400 µg/mL IdU for 45 minutes. Cells were then washed and released into medium containing 400 µg/mL CldU for 30 minutes before arrest of the culture by addition of NaN3 to a final concentration of 0.1%. 1.2x10^8^ cells were washed in ice cold TE50 and cell pellets frozen in liquid nitrogen and stored at -80°C. Cell cycle stage of samples as well as arrest and release efficiency was determined by PI staining and flow cytometry (see above).

Cells were washed twice in ice cold SP1 buffer (1.2 M sorbitol, 50 mM NaCitrate, 40 mM EDTA) and resuspended in 150 µl SP1 buffer + 0.8 mg/mL zymolase (MilliPore), 1 mg/mL lysing enzyme (Sigma), 1% β-mercaptoethanol. The cell suspension was mixed with 170 µl 1.2% low-melting point agarose (Sigma), distributed into plug molds (BioRad) and allowed to set at 4°C. Plugs were incubated 4-6h at 37°C in SP1 buffer + 0.2 mg/mL zymolase, 0.25 mg/mL lysing enzyme, 1% β- marcaptoethanol before being washed in 0.5 M EDTA and incubated overnight in 20 mg/mL proteinase K (Sigma), 400 mM EDTA, 10% sarkosyl. Plugs were then washed thrice in TE buffer and stored at 4°C for at least three days.

One plug per sample was melted at 68°C in 1 mL 100 mM MES buffer pH 6.5 prior to addition of β-agarase (New England Biolabs) and incubation overnight at 42°C. The DNA solution was combed onto silanized CombiCoverslips (GenomicVision) using the GenomicVision molecular combing system. Slides were baked 4h at 60°C before denaturation of the DNA for 10 min in 0.5 M NaOH, washing in PBS and sequential 5 min dehydrations in 70%, 80% and 96% ethanol. Slides were blocked 25 minutes at 37°C in the dark in PBS + 0.05% Tween-20, 1.5% BSA prior to 1 hour incubation in mouse anti-IdU (1:5, 347580, BD Bioscience) and rat anti-CldU-FITC (1:25, Ab74545, Abcam) followed by washes in PBS + 0.05% Tween-20 and 30 minutes incubation in goat anti-mouse Alexa fluor 633 and goat anti-rat Alexa fluor 488 (1:25, A21050 and A11006, Molecular Probes). CombiCoverslips were mounted with ProLong Diamond Antifade reagent (P36965, Molecular Probes) and allowed to set overnight at room temperature in the dark.

DNA fibres were imaged on a Zeiss confocal LSM700 microscope. Length of IdU and CldU tracts were measured in Fiji (Schindelin *et al*., 2012) and converted to kilo bases based on the pixel-to-µm conversion factor of the microscope (1 px = 0.13 µm for 746 x 746 px images) and the constant stretching factor of the GenomicVision molecular combing system (1 µm = 2 kb). Representations of data in graphs were done in Prism (GraphPad software, Inc.) and statistically significant differences in IdU and CldU tract lengths observed between cell populations were determined by Mann-Whitney test, while differences in fork restart were determined by Fisher’s exact test.

### Data availability

The mass spectrometry proteomics data have been deposited to the ProteomeXchange Consortium via the PRIDE (Perez-Riverol *et al*, 2019) partner repository with the dataset identifier PXD028478.

Reviewer account details:

**PRIDE website**: http://www.ebi.ac.uk/pride

**Password:** KbIyVCKe

## References

Abbas T, Keaton MA, Dutta A (2013) Genomic instability in cancer. Cold Spring Harbor perspectives in biology 5: a012914

Alcasabas AA, Osborn AJ, Bachant J, Hu F, Werler PJ, Bousset K, Furuya K, Diffley JF, Carr AM, Elledge SJ (2001) Mrc1 transduces signals of DNA replication stress to activate Rad53. Nat Cell Biol 3: 958–965

Andress EJ, Holic R, Edelmann MJ, Kessler BM, Yu VP (2011) Dia2 controls transcription by mediating assembly of the RSC complex. PLoS One 6: e21172

Appanah R, Lones EC, Aiello U, Libri D, De Piccoli G (2020) Sen1 Is Recruited to Replication Forks via Ctf4 and Mrc1 and Promotes Genome Stability. Cell reports 30: 2094–2105 e2099

Baek GH, Cheng H, Choe V, Bao X, Shao J, Luo S, Rao H (2013) Cdc48: a swiss army knife of cell biology. J Amino Acids 2013: 183421

Baek GH, Kim I, Rao H (2011) The Cdc48 ATPase modulates the interaction between two proteolytic factors Ufd2 and Rad23. Proc Natl Acad Sci U S A 108: 13558–13563

Bando M, Katou Y, Komata M, Tanaka H, Itoh T, Sutani T, Shirahige K (2009) Csm3, Tof1, and Mrc1 form a heterotrimeric mediator complex that associates with DNA replication forks. J Biol Chem 284: 34355–34365

Baretic D, Jenkyn-Bedford M, Aria V, Cannone G, Skehel M, Yeeles JTP (2020) Cryo-EM Structure of the Fork Protection Complex Bound to CMG at a Replication Fork. Mol Cell 78: 926–940 e913

Bays NW, Wilhovsky SK, Goradia A, Hodgkiss-Harlow K, Hampton RY (2001) HRD4/NPL4 is required for the proteasomal processing of ubiquitinated ER proteins. Mol Biol Cell 12: 4114–4128

Bergink S, Ammon T, Kern M, Schermelleh L, Leonhardt H, Jentsch S (2013) Role of Cdc48/p97 as a SUMO-targeted segregase curbing Rad51-Rad52 interaction. Nat Cell Biol 15: 526–532

Blastyak A, Pinter L, Unk I, Prakash L, Prakash S, Haracska L (2007) Yeast Rad5 protein required for postreplication repair has a DNA helicase activity specific for replication fork regression. Mol Cell 28: 167–175

Brangwynne Clifford P, Tompa P, Pappu Rohit V (2015) Polymer physics of intracellular phase transitions. Nature Physics 11: 899–904

Chaudhury I, Koepp DM (2017) Degradation of Mrc1 promotes recombination-mediated restart of stalled replication forks. Nucleic Acids Res 45: 2558–2570

Chen SH, Zhou H (2009) Reconstitution of Rad53 activation by Mec1 through adaptor protein Mrc1. J Biol Chem 284: 18593–18604

Chini CC, Chen J (2003) Human claspin is required for replication checkpoint control. J Biol Chem 278: 30057–30062

Chini CC, Chen J (2004) Claspin, a regulator of Chk1 in DNA replication stress pathway. DNA Repair (Amst) 3: 1033–1037

Chini CC, Chen J (2006) Repeated phosphopeptide motifs in human Claspin are phosphorylated by Chk1 and mediate Claspin function. J Biol Chem 281: 33276–33282

Cho RJ, Campbell MJ, Winzeler EA, Steinmetz L, Conway A, Wodicka L, Wolfsberg TG, Gabrielian AE, Landsman D, Lockhart DJ et al (1998) A genome-wide transcriptional analysis of the mitotic cell cycle. Mol Cell 2: 65–73

Dai RM, Chen E, Longo DL, Gorbea CM, Li CC (1998) Involvement of valosin-containing protein, an ATPase Co-purified with IkappaBalpha and 26 S proteasome, in ubiquitin-proteasome-mediated degradation of IkappaBalpha. J Biol Chem 273: 3562–3573

Dai RM, Li CC (2001) Valosin-containing protein is a multi-ubiquitin chain-targeting factor required in ubiquitin-proteasome degradation. Nat Cell Biol 3: 740–744

de Lichtenberg U, Jensen TS, Jensen LJ, Brunak S (2003) Protein feature based identification of cell cycle regulated proteins in yeast. J Mol Biol 329: 663–674

den Brave F, Cairo LV, Jagadeesan C, Ruger-Herreros C, Mogk A, Bukau B, Jentsch S (2020) Chaperone-Mediated Protein Disaggregation Triggers Proteolytic Clearance of Intra-nuclear Protein Inclusions. Cell reports 31: 107680

Duch A, Canal B, Barroso SI, Garcia-Rubio M, Seisenbacher G, Aguilera A, de Nadal E, Posas F (2018) Multiple signaling kinases target Mrc1 to prevent genomic instability triggered by transcription-replication conflicts. Nature communications 9: 379

Elsasser S, Finley D (2005) Delivery of ubiquitinated substrates to protein-unfolding machines. Nat Cell Biol 7: 742–749

Erdeniz N, Mortensen UH, Rothstein R (1997) Cloning-free PCR-based allele replacement methods. Genome Res 7: 1174–1183

Fong CM, Arumugam A, Koepp DM (2013) The Saccharomyces cerevisiae F-box protein Dia2 is a mediator of S-phase checkpoint recovery from DNA damage. Genetics 193: 483–499

Franz A, Orth M, Pirson PA, Sonneville R, Blow JJ, Gartner A, Stemmann O, Hoppe T (2011) CDC-48/p97 coordinates CDT-1 degradation with GINS chromatin dissociation to ensure faithful DNA replication. Mol Cell 44: 85–96

Frattini C, Villa-Hernandez S, Pellicano G, Jossen R, Katou Y, Shirahige K, Bermejo R (2017) Cohesin Ubiquitylation and Mobilization Facilitate Stalled Replication Fork Dynamics. Mol Cell 68: 758–772 e754

Gallagher PS, Clowes Candadai SV, Gardner RG (2014) The requirement for Cdc48/p97 in nuclear protein quality control degradation depends on the substrate and correlates with substrate insolubility. J Cell Sci 127: 1980–1991

Gallina I, Colding C, Henriksen P, Beli P, Nakamura K, Offman J, Mathiasen DP, Silva S, Hoffmann E, Groth A et al (2015) Cmr1/WDR76 defines a nuclear genotoxic stress body linking genome integrity and protein quality control. Nature communications 6: 6533

Gardner RG, Nelson ZW, Gottschling DE (2005) Degradation-mediated protein quality control in the nucleus. Cell 120: 803–815

Gasch AP, Huang M, Metzner S, Botstein D, Elledge SJ, Brown PO (2001) Genomic expression responses to DNA-damaging agents and the regulatory role of the yeast ATR homolog Mec1p. Mol Biol Cell 12: 2987–3003.

Gellon L, Kaushal S, Cebrian J, Lahiri M, Mirkin SM, Freudenreich CH (2019) Mrc1 and Tof1 prevent fragility and instability at long CAG repeats by their fork stabilizing function. Nucleic Acids Res 47: 794–805

Gispan A, Carmi M, Barkai N (2014) Checkpoint-independent scaling of the Saccharomyces cerevisiae DNA replication program. BMC Biol 12: 79

Hartmann-Petersen R, Wallace M, Hofmann K, Koch G, Johnsen AH, Hendil KB, Gordon C (2004) The Ubx2 and Ubx3 cofactors direct Cdc48 activity to proteolytic and nonproteolytic ubiquitin-dependent processes. Curr Biol 14: 824–828

Haslbeck M, Braun N, Stromer T, Richter B, Model N, Weinkauf S, Buchner J (2004) Hsp42 is the general small heat shock protein in the cytosol of Saccharomyces cerevisiae. EMBO J 23: 638–649

Hodgson B, Calzada A, Labib K (2007) Mrc1 and Tof1 regulate DNA replication forks in different ways during normal S phase. Mol Biol Cell 18: 3894–3902

Janke C, Magiera MM, Rathfelder N, Taxis C, Reber S, Maekawa H, Moreno-Borchart A, Doenges G, Schwob E, Schiebel E et al (2004) A versatile toolbox for PCR-based tagging of yeast genes: new fluorescent proteins, more markers and promoter substitution cassettes. Yeast 21: 947–962

Jentsch S, Psakhye I (2013) Control of nuclear activities by substrate-selective and protein-group SUMOylation. Annu Rev Genet 47: 167–186

Jentsch S, Rumpf S (2007) Cdc48 (p97): a "molecular gearbox" in the ubiquitin pathway? Trends Biochem Sci 32: 6–11

Katou Y, Kanoh Y, Bando M, Noguchi H, Tanaka H, Ashikari T, Sugimoto K, Shirahige K (2003) S-phase checkpoint proteins Tof1 and Mrc1 form a stable replication-pausing complex. Nature 424: 1078–1083

Khmelinskii A, Keller PJ, Bartosik A, Meurer M, Barry JD, Mardin BR, Kaufmann A, Trautmann S, Wachsmuth M, Pereira G et al (2012) Tandem fluorescent protein timers for in vivo analysis of protein dynamics. Nat Biotechnol 30: 708–714

Kile AC, Koepp DM (2010) Activation of the S-phase checkpoint inhibits degradation of the F-box protein Dia2. Mol Cell Biol 30: 160–171

Labib K, De Piccoli G (2011) Surviving chromosome replication: the many roles of the S-phase checkpoint pathway. Philos Trans R Soc Lond B Biol Sci 366: 3554–3561

Lazar C, Gatto L, Ferro M, Bruley C, Burger T (2016) Accounting for the Multiple Natures of Missing Values in Label-Free Quantitative Proteomics Data Sets to Compare Imputation Strategies. J Proteome Res 15: 1116–1125

Lisby M, Rothstein R, Mortensen UH (2001) Rad52 forms DNA repair and recombination centers during S phase. Proc Natl Acad Sci USA 98: 8276–8282

Lou H, Komata M, Katou Y, Guan Z, Reis CC, Budd M, Shirahige K, Campbell JL (2008) Mrc1 and DNA polymerase epsilon function together in linking DNA replication and the S phase checkpoint. Mol Cell 32: 106–117

Luke B, Versini G, Jaquenoud M, Zaidi IW, Kurz T, Pintard L, Pasero P, Peter M (2006) The cullin Rtt101p promotes replication fork progression through damaged DNA and natural pause sites. Curr Biol 16: 786–792

Maculins T, Nkosi PJ, Nishikawa H, Labib K (2015) Tethering of SCF to the Replisome Promotes Efficient Ubiquitylation and Disassembly of the CMG Helicase. *Curr Biol*

Mailand N, Bekker-Jensen S, Bartek J, Lukas J (2006) Destruction of Claspin by SCFbetaTrCP restrains Chk1 activation and facilitates recovery from genotoxic stress. Mol Cell 23: 307–318

Maric M, Maculins T, De Piccoli G, Labib K (2014) Cdc48 and a ubiquitin ligase drive disassembly of the CMG helicase at the end of DNA replication. Science 346: 1253596

Maric M, Mukherjee P, Tatham MH, Hay R, Labib K (2017) Ufd1-Npl4 Recruit Cdc48 for Disassembly of Ubiquitylated CMG Helicase at the End of Chromosome Replication. Cell reports 18: 3033–3042

Martin Y, Cabrera E, Amoedo H, Hernandez-Perez S, Dominguez-Kelly R, Freire R (2015) USP29 controls the stability of checkpoint adaptor Claspin by deubiquitination. Oncogene 34: 1058–1063

McClure AW, Diffley JF (2021) Rad53 checkpoint kinase regulation of DNA replication fork rate via Mrc1 phosphorylation. eLife 10

McGarry E, Gaboriau D, Rainey M, Restuccia U, Bachi A, Santocanale C (2016) The deubiquitinase USP9X maintains DNA replication fork stability and DNA damage checkpoint responses by regulating CLASPIN during S-phase. Cancer Res

Meier F, Brunner AD, Koch S, Koch H, Lubeck M, Krause M, Goedecke N, Decker J, Kosinski T, Park MA et al (2018) Online Parallel Accumulation-Serial Fragmentation (PASEF) with a Novel Trapped Ion Mobility Mass Spectrometer. Mol Cell Proteomics 17: 2534–2545

Menin L, Ursich S, Trovesi C, Zellweger R, Lopes M, Longhese MP, Clerici M (2018) Tel1/ATM prevents degradation of replication forks that reverse after topoisomerase poisoning. EMBO Rep 19

Meszaros B, Erdos G, Dosztanyi Z (2018) IUPred2A: context-dependent prediction of protein disorder as a function of redox state and protein binding. Nucleic Acids Res 46: W329–W337

Meyer HH, Wang Y, Warren G (2002) Direct binding of ubiquitin conjugates by the mammalian p97 adaptor complexes, p47 and Ufd1-Npl4. EMBO J 21: 5645–5652

Miller SB, Ho CT, Winkler J, Khokhrina M, Neuner A, Mohamed MY, Guilbride DL, Richter K, Lisby M, Schiebel E et al (2015) Compartment-specific aggregases direct distinct nuclear and cytoplasmic aggregate deposition. EMBO J 34: 778–797

Mimura S, Komata M, Kishi T, Shirahige K, Kamura T (2009) SCF(Dia2) regulates DNA replication forks during S-phase in budding yeast. EMBO J 28: 3693–3705

Morawska M, Ulrich HD (2013) An expanded tool kit for the auxin-inducible degron system in budding yeast. Yeast 30: 341–351

Moreno SP, Bailey R, Campion N, Herron S, Gambus A (2014) Polyubiquitylation drives replisome disassembly at the termination of DNA replication. Science 346: 477–481

Morohashi H, Maculins T, Labib K (2009) The amino-terminal TPR domain of Dia2 tethers SCF(Dia2) to the replisome progression complex. Curr Biol 19: 1943–1949

Naylor ML, Li JM, Osborn AJ, Elledge SJ (2009) Mrc1 phosphorylation in response to DNA replication stress is required for Mec1 accumulation at the stalled fork. Proc Natl Acad Sci U S A 106: 12765–12770

Nie M, Aslanian A, Prudden J, Heideker J, Vashisht AA, Wohlschlegel JA, Yates JR, 3rd, Boddy MN (2012) Dual recruitment of Cdc48 (p97)-Ufd1-Npl4 ubiquitin-selective segregase by small ubiquitin-like modifier protein (SUMO) and ubiquitin in SUMO-targeted ubiquitin ligase-mediated genome stability functions. J Biol Chem 287: 29610–29619

Nishimura K, Fukagawa T, Takisawa H, Kakimoto T, Kanemaki M (2009) An auxin-based degron system for the rapid depletion of proteins in nonplant cells. Nature methods 6: 917–922

O’Neill BM, Szyjka SJ, Lis ET, Bailey AO, Yates JR, 3rd, Aparicio OM, Romesberg FE (2007) Pph3-Psy2 is a phosphatase complex required for Rad53 dephosphorylation and replication fork restart during recovery from DNA damage. Proc Natl Acad Sci U S A 104: 9290–9295

On KF, Beuron F, Frith D, Snijders AP, Morris EP, Diffley JF (2014) Prereplicative complexes assembled in vitro support origin-dependent and independent DNA replication. EMBO J 33: 605–620

Osborn AJ, Elledge SJ (2003) Mrc1 is a replication fork component whose phosphorylation in response to DNA replication stress activates Rad53. Genes Dev 17: 1755–1767.

Perez-Riverol Y, Csordas A, Bai J, Bernal-Llinares M, Hewapathirana S, Kundu DJ, Inuganti A, Griss J, Mayer G, Eisenacher M et al (2019) The PRIDE database and related tools and resources in 2019: improving support for quantification data. Nucleic Acids Res 47: D442–D450

Peschiaroli A, Dorrello NV, Guardavaccaro D, Venere M, Halazonetis T, Sherman NE, Pagano M (2006) SCFbetaTrCP-mediated degradation of Claspin regulates recovery from the DNA replication checkpoint response. Mol Cell 23: 319–329

Petermann E, Helleday T, Caldecott KW (2008) Claspin promotes normal replication fork rates in human cells. Mol Biol Cell 19: 2373–2378

Pramila T, Miles S, GuhaThakurta D, Jemiolo D, Breeden LL (2002) Conserved homeodomain proteins interact with MADS box protein Mcm1 to restrict ECB-dependent transcription to the M/G1 phase of the cell cycle. Genes Dev 16: 3034–3045

Pramila T, Wu W, Miles S, Noble WS, Breeden LL (2006) The Forkhead transcription factor Hcm1 regulates chromosome segregation genes and fills the S-phase gap in the transcriptional circuitry of the cell cycle. Genes Dev 20: 2266–2278

Psakhye I, Castellucci F, Branzei D (2019) SUMO-Chain-Regulated Proteasomal Degradation Timing Exemplified in DNA Replication Initiation. Mol Cell 76: 632–645 e636

Psakhye I, Jentsch S (2012) Protein group modification and synergy in the SUMO pathway as exemplified in DNA repair. Cell 151: 807–820

Ramadan K, Halder S, Wiseman K, Vaz B (2017) Strategic role of the ubiquitin-dependent segregase p97 (VCP or Cdc48) in DNA replication. Chromosoma 126: 17–32

Raman M, Havens CG, Walter JC, Harper JW (2011) A genome-wide screen identifies p97 as an essential regulator of DNA damage-dependent CDT1 destruction. Mol Cell 44: 72–84

Rape M, Hoppe T, Gorr I, Kalocay M, Richly H, Jentsch S (2001) Mobilization of processed, membrane-tethered SPT23 transcription factor by CDC48(UFD1/NPL4), a ubiquitin-selective chaperone. Cell 107: 667–677

Reid R, Lisby M, Rothstein R (2002) Cloning-free genome alterations in *Saccharomyce cerevisiae* using adaptamer-mediated PCR. Methods Enzymol 350: 258–277

Reusswig K-U, Bittmann J, Peritore M, Wierer M, Mann M, Pfander B (2021) Unscheduled DNA replication in G1 causes genome instability through head-to-tail replication fork collisions. bioRxiv: 2021.2009.2006.459115

Rumpf S, Jentsch S (2006) Functional division of substrate processing cofactors of the ubiquitin-selective Cdc48 chaperone. Mol Cell 21: 261–269

Santos A, Colaço AR, Nielsen AB, Niu L, Geyer PE, Coscia F, Albrechtsen NJW, Mundt F, Jensen LJ, Mann M (2020) Clinical Knowledge Graph Integrates Proteomics Data into Clinical Decision-Making. bioRxiv: 2020.2005.2009.084897

Schindelin J, Arganda-Carreras I, Frise E, Kaynig V, Longair M, Pietzsch T, Preibisch S, Rueden C, Saalfeld S, Schmid B et al (2012) Fiji: an open-source platform for biological-image analysis. Nat Methods 9: 676–682

Schmidt KH, Kolodner RD (2006) Suppression of spontaneous genome rearrangements in yeast DNA helicase mutants. Proc Natl Acad Sci U S A 103: 18196–18201

Sherman F, Fink, G. R. & Hicks, J. B. (1986) Methods in Yeast Genetics. Cold Spring Harbor Laboratory, Cold Spring Harbor, NY

Shyu YJ, Hu CD (2008) Fluorescence complementation: an emerging tool for biological research. Trends in biotechnology 26: 622–630

Silva S, Altmannova, V., Luke-Glaser, S., Henriksen, P., Gallina, I., Yang, X., Choudhary, C., Luke, B., Krejci, L. and Lisby, M (2016) Mte1 interacts with Mph1 and promotes crossover recombination and telomere maintenance. Genes & Development in press

Silva S, Gallina I, Eckert-Boulet N, Lisby M (2012) Live Cell Microscopy of DNA Damage Response in *Saccharomyces cerevisiae*. Methods Mol Biol 920: 433–443

Spellman PT, Sherlock G, Zhang MQ, Iyer VR, Anders K, Eisen MB, Brown PO, Botstein D, Futcher B (1998) Comprehensive identification of cell cycle-regulated genes of the yeast Saccharomyces cerevisiae by microarray hybridization. Mol Biol Cell 9: 3273–3297

Stein A, Ruggiano A, Carvalho P, Rapoport TA (2014) Key steps in ERAD of luminal ER proteins reconstituted with purified components. Cell 158: 1375–1388

Sung MK, Lim G, Yi DG, Chang YJ, Yang EB, Lee K, Huh WK (2013) Genome-wide bimolecular fluorescence complementation analysis of SUMO interactome in yeast. Genome Res 23: 736–746

Szyjka SJ, Viggiani CJ, Aparicio OM (2005) Mrc1 is required for normal progression of replication forks throughout chromatin in S. cerevisiae. Mol Cell 19: 691–697

Tourriere H, Versini G, Cordon-Preciado V, Alabert C, Pasero P (2005) Mrc1 and Tof1 promote replication fork progression and recovery independently of Rad53. Mol Cell 19: 699–706

Uzunova SD, Zarkov AS, Ivanova AM, Stoynov SS, Nedelcheva-Veleva MN (2014) The subunits of the S-phase checkpoint complex Mrc1/Tof1/Csm3: dynamics and interdependence. Cell division 9: 4

Verma R, Chen S, Feldman R, Schieltz D, Yates J, Dohmen J, Deshaies RJ (2000) Proteasomal proteomics: identification of nucleotide-sensitive proteasome-interacting proteins by mass spectrometric analysis of affinity-purified proteasomes. Mol Biol Cell 11: 3425–3439

Verma R, Oania R, Fang R, Smith GT, Deshaies RJ (2011) Cdc48/p97 mediates UV-dependent turnover of RNA Pol II. Mol Cell 41: 82–92

Voordeckers K, Colding C, Grasso L, Pardo B, Hoes L, Kominek J, Gielens K, Dekoster K, Gordon J, Van der Zande E et al (2020) Ethanol exposure increases mutation rate through error-prone polymerases. Nature communications 11: 3664

Xu H, Boone C, Klein HL (2004) Mrc1 is required for sister chromatid cohesion to aid in recombination repair of spontaneous damage. Mol Cell Biol 24: 7082–7090

Yamauchi Y, Izawa S (2016) Prioritized Expression of BTN2 of Saccharomyces cerevisiae under Pronounced Translation Repression Induced by Severe Ethanol Stress. Front Microbiol 7: 1319

Ye Y, Meyer HH, Rapoport TA (2003) Function of the p97-Ufd1-Npl4 complex in retrotranslocation from the ER to the cytosol: dual recognition of nonubiquitinated polypeptide segments and polyubiquitin chains. J Cell Biol 162: 71–84

Yeeles JTP, Janska A, Early A, Diffley JFX (2017) How the Eukaryotic Replisome Achieves Rapid and Efficient DNA Replication. Mol Cell 65: 105–116

Zaidi IW, Rabut G, Poveda A, Scheel H, Malmstrom J, Ulrich H, Hofmann K, Pasero P, Peter M, Luke B (2008) Rtt101 and Mms1 in budding yeast form a CUL4(DDB1)- like ubiquitin ligase that promotes replication through damaged DNA. EMBO Rep 9: 1034–1040

Zhang J, Shi D, Li X, Ding L, Tang J, Liu C, Shirahige K, Cao Q, Lou H (2017) Rtt101- Mms1-Mms22 coordinates replication-coupled sister chromatid cohesion and nucleosome assembly. EMBO Rep 18: 1294–1305

Zhao X, Wu CY, Blobel G (2004) Mlp-dependent anchorage and stabilization of a desumoylating enzyme is required to prevent clonal lethality. J Cell Biol 167: 605–611

